# Chloroplast cold-resistance is mediated by the acidic domain of the RNA binding protein CP31A

**DOI:** 10.1101/832337

**Authors:** Ayako Okuzaki, Marie-Kristin Lehniger, Jose M Muino, Benjamin Lenzen, Thilo Rühe, Dario Leister, Uwe Ohler, Christian Schmitz-Linneweber

**Author notes:** College of Agriculture, Tamagawa University, Tamagawagakuen 6-1-1, Machida-shi, Tokyo, Japan (Ayako Okuzaki). equally contributed (Ayako Okuzaki, Marie-Kristin Lehniger). corresponding author: Christian Schmitz-Linneweber. Author contributions C.S.-L. conceived the original research plan; D.L. supervised the experiments; A.O. generated the transgenic plant lines and performed their analysis. M.-K. L. performed the RIP-Seq and microarray experiments; J.M.M. analyzed the RIP-Seq data. B.L. performed the SSMART analysis; T.R. performed mutant analyses. C.S.-L., D.L. and U.O. supervised the project, designed the experiments and analyzed the data; C.S.-L. wrote the article with contributions of all the authors; C.S.-L. agrees to serve as the author responsible for contact and ensures communication.

## Abstract

Chloroplast RNA metabolism is characterized by long-lived mRNAs that undergo a multitude of post-transcriptional processing events. Chloroplast RNA accumulation responds to environmental cues, foremost light and temperature. A large number of nuclear-encoded RNA-binding proteins (RBPs) are required for chloroplast RNA metabolism, but we do not yet know how chloroplast RBPs convert abiotic signals into gene expression changes. Previous studies showed that the chloroplast ribonucleoprotein 31A (CP31A) is required for the stabilization of multiple chloroplast mRNAs in the cold, and that the phosphorylation of CP31A at various residues within its N-terminal acidic domain (AD) can alter its affinity for RNA *in vitro*. Loss of CP31A leads to cold sensitive plants that exhibit bleached tissue at the center of the vegetative rosette. Here, by applying RIP-Seq, we demonstrated that CP31A shows increased affinity for a large number of chloroplast RNAs *in vivo* in the cold. Among the main targets of CP31A were RNAs encoding subunits of the NDH complex and loss of CP31A lead to reduced accumulation of *ndh* transcripts. Deletion analyses revealed that cold-dependent RNA binding and cold resistance of chloroplast development both depend on the AD of CP31A. Together, our analysis established the AD of CP31A as a key mediator of cold acclimation of the chloroplast transcriptome.

**One sentence summary:** Cold exposure induces increased RNA association of the RRM protein CP31A, which mediates cold-resistance of *Arabidopsis thaliana* via its acidic domain

## Introduction

Chloroplasts contain genetic information that is essential for photosynthesis. The expression of this information is realized by a unique mixture of ancestral bacterial and derived eukaryotic features (Barkan, 2011). Chloroplast gene expression adapts to various environmental changes, including light and temperature (e.g. Klein, 1991; Mentzen and Wurtele, 2008; Cho et al., 2009; Castandet et al., 2016). Contributions to such acclimation processes have been described on the transcriptional level (Pfannschmidt, 2003; Tsunoyama et al., 2004), but post-transcriptional processes are likely to dominate (Deng and Gruissem, 1987; Eberhard et al., 2002; Udy et al., 2012). One key change in posttranscriptional processes between chloroplast and their bacterial ancestors are the vastly increased RNA half lives in the organelle. In bacteria, transcription and translation are usually directly coupled and mRNAs have short half-lives (in the range of minutes; Selinger et al., 2003). In chloroplasts, the half-lives of mRNAs are long (in the range of hours) and untranslated RNAs accumulate in large amounts (Klaff and Gruissem, 1991; Germain et al., 2012). The turnover rates of chloroplast RNAs change in response to developmental and environmental cues, which is suggesting that RNA stability is regulated in chloroplasts (Deng et al., 1989; Klaff and Gruissem, 1991; Biehl et al., 2005; Bollenbach et al., 2007; Germain et al., 2013; Manavski et al., 2018), but the underlying regulatory factors are largely unknown. Possible candidates for regulators of RNA stability are pentatricopeptide repeat (PPR) proteins and chloroplast ribonucleoproteins (cpRNPs). PPR proteins specifically associate with one or few RNAs (Barkan and Small, 2014), while cpRNPs are generalists that bind to a large number of mRNAs (Kupsch et al., 2012; Teubner et al., 2017). Two cpRNPs named CP29A and CP31A were previously shown to be required for the accumulation of chloroplast RNAs in the cold, which makes them interesting candidates to be mediators of global RNA stability during the acclimation to changing temperature conditions (Kupsch et al., 2012).

The cpRNP protein family consists of ten members in *Arabidopsis thaliana* (Ruwe et al., 2011). All cpRNPs are targeted to the chloroplast post-translationally, and dedicated import receptors appear to be responsible for their transport across the chloroplast envelope (Li and Sugiura, 1990; Grimmer et al., 2014). cpRNPs are highly regulated proteins that react to various external and internal signals, particularly light, which controls both their expression and their protein modification state (summarized in Ruwe et al., 2011). Several cpRNPs have been identified as phosphoproteins (Reiland et al., 2009) and an N-terminally acetylated isoform of CP29A was shown to respond rapidly to changes in light and developmental stages (Wang et al., 2006). The phosphorylation of cpRNPs can alter their RNA-binding characteristics *in vitro* (Lisitsky and Schuster, 1995; Loza-Tavera et al., 2006), but evidence for condition-dependent RNA binding of cpRNPs or any other chloroplast RBPs *in vivo* are lacking.

Several molecular functions have been suggested for cpRNPs. *In vitro*, a tobacco homolog of *Arabidopsis* CP31A has been shown to support RNA editing of multiple sites (Hirose and Sugiura, 2001). Other cpRNPs have been reported to be required for the 3’-end processing of several mRNAs (Schuster and Gruissem, 1991; Hayes et al., 1996; Schuster et al., 1999). They also support the ribozymatic maturation of a viroid RNA genome (Daros and Flores, 2002). The most notable and general function of cpRNPs is however their role in stabilizing mRNAs. *In vitro*, cpRNPs protect the mRNA encoding the D1 subunit of photosystem II against degradation (Nakamura et al., 2001). *In vivo*, they are required for the accumulation of a multitude of mRNAs (Kupsch et al., 2012; Teubner et al., 2017). Co-immunoprecipitation analyses have demonstrated that cpRNPs are associated with multiple chloroplast RNAs (Kupsch et al., 2012; Teubner et al., 2017), and that they prefer unprocessed (unspliced) RNAs over mature forms (Nakamura et al., 1999). Together, the data obtained from these functional analyses show that cpRNPs have a broad target range and contribute to various RNA-processing events and to RNA stabilization.

All cpRNPs share a similar design with two RNA recognition motifs (RRMs) that are preceded by a domain rich in acidic residues (acidic domain; AD). While the RRMs are well characterized RNA binding domains (Li and Sugiura, 1991; Ye and Sugiura, 1992; Lisitsky et al., 1995), the role of the AD remains to be determined. The cpRNP CP31A stands out from the family in having a particularly long acidic domain, which contains two phosphorylated serine residues within a short repeat element of five amino acids (Reiland et al., 2009). CP31A’s AD was suggested to be important for RNA editing based on *in vitro* assays (Hirose and Sugiura, 2001). The AD of the spinach homolog of CP31A is supportive of RNA binding *in vitro*, but does not bind RNA itself (Lisitsky and Schuster, 1995). Recently, plastid casein kinase II (pCKII) was demonstrated to phosphorylate the acidic domain of CP31A *in vitro* (Schonberg et al., 2014). Genetic analyses have demonstrated that *Arabidopsis* CP31A supports RNA editing at multiple sites and modulates the stability of multiple mRNAs (Tillich et al., 2009; Kupsch et al., 2012). An RNA strongly affected in *cp31a* mutants was the *ndhF* mRNA, which encodes a subunit of the NDH complex. CP31A binds to the 3’-UTR of *ndhF* and is required for the generation of the *ndhF* 3’-terminus (Kupsch et al., 2012). Mutants of CP31A display cold sensitivity: the germination rate of null mutants is reduced and their newly emerging leaf tissue bleaches at 8°C (Kupsch et al., 2012). All analyzed proteins of the photosynthetic apparatus are reduced in this defective tissue, which is at least in part due to multiple defects in RNA processing (e.g., RNA splicing, RNA editing, and intercistronic processing), but is likely primarily caused by the strong reduction of multiple chloroplast mRNAs (Kupsch et al., 2012). However, it remains unclear how CP31A mechanistically confers cold resistance and whether it directly perceives cold as a signal. We hypothesized that the AD could act as a regulatory domain for RNA binding and cold-responsiveness. We therefore analyzed the ability of CP31A to associate with RNA in response to cold and assessed the role of the AD for CP31A’s ability to bind and process chloroplast RNA. Our results demonstrate that the AD of CP31A is essential for plant cold acclimation, at least in part because of its supportive role for cold-dependent RNA binding.

## Results

### The RRM domains of CP31A are sufficient for RNA editing of CP31A-dependent sites

The binding of RBPs to their RNA targets can be altered by protein modifications. For several members of the cpRNP family, phosphorylation has been demonstrated to alter RNA binding *in vitro* (Lisitsky and Schuster, 1995; Loza-Tavera et al., 2006; Reiland et al., 2009). As phosphoproteomic analyses identified several phosphorylation sites within the acidic N-terminal domain of CP31A (Reiland et al., 2009; Schonberg et al., 2014), we decided to test the function of this acidic phosphodomain by deletion mutagenesis. We designed three T-DNA based constructs to express CP31A variants in *Arabidopsis* (Fig. 1A). The first two expressed variants lack the AD. One was driven by the 35S promoter, while the other was driven by the endogenous genomic region upstream of the transcriptional start site, presumably encompassing the unknown *CP31A* promoter (12769646 to 12767953 of the *Arabidopsis thaliana* chromosome 4 sequence; nucleotides –1 to –1694 relative to the *CP31A* start codon). These constructs were introduced into a *cp31a*-deficient background (homozygous *cp31a*-1 null allele) by *Agrobacterium*-mediated gene transfer. In addition, the full-length protein driven by its native promoter was used as a positive control. All plant lines retrieved showed normal development and were indistinguishable from wt plants under normal growth conditions (see below).

**Figure 1:**
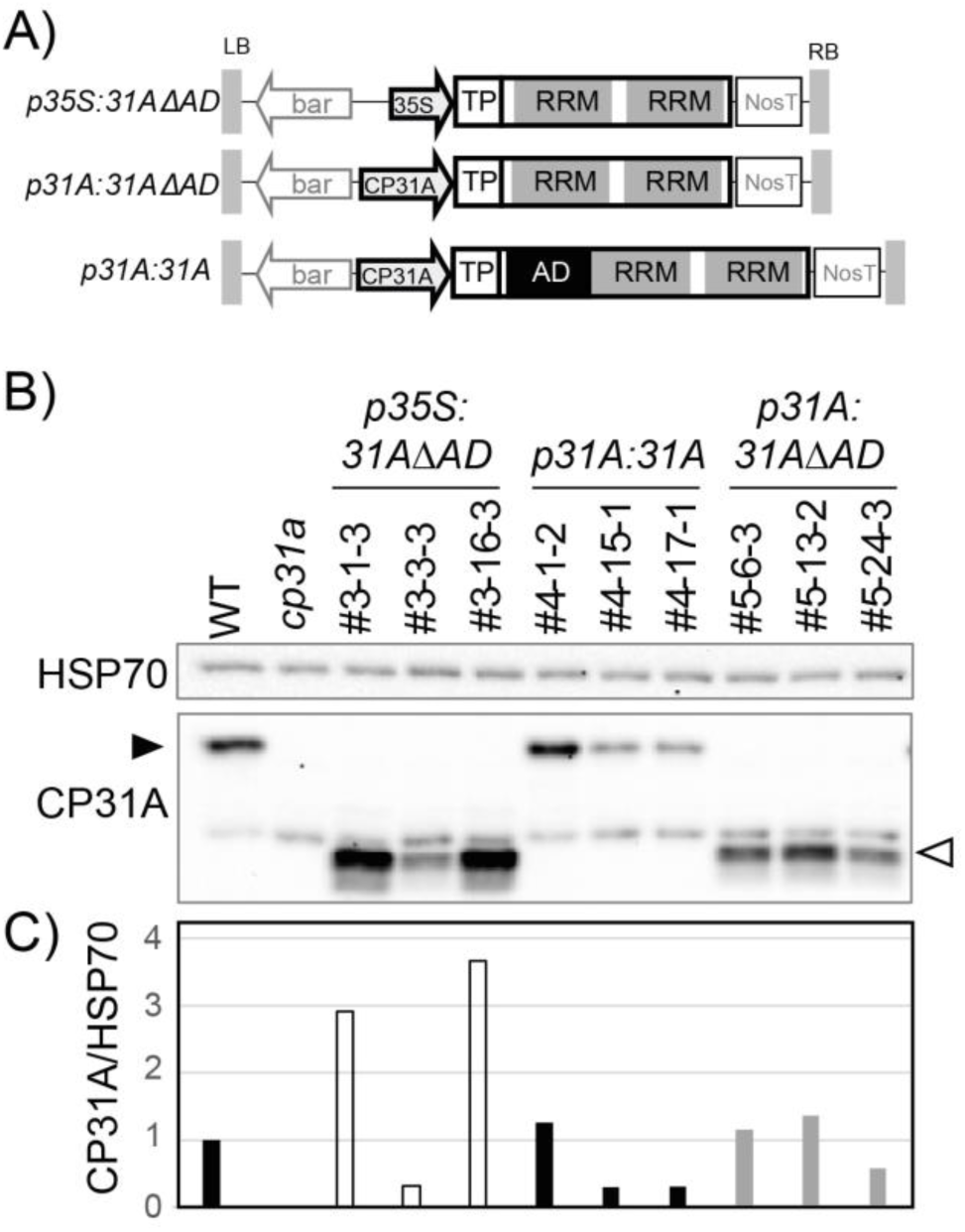
Construction of AD deletion mutants. A) Schematic overview of constructs used to complement *cp31a* null mutants. bar = selectable marker; black arrows indicate promoters. TP = transit peptide. NosT = the terminator of the nos gene. LB/RB = borders of the *Agrobacterium* T-DNA. B) Western analysis of transgenic lines. Blots were probed with a HSP70 antibody to control for loading and with an antiserum recognizing CP31A. The position of the full-length protein and the truncated, AD-less version are indicated by filled and open arrowheads, respectively. C) Quantification of CP31A signals normalized to HSP70 signals (from (B)).

We assessed CP31A protein accumulation in our transgenic lines by using an antibody that reacts to two peptides: one situated in the linker between the two RRM domains and another at the C-terminus of the protein (Kupsch et al. 2012). The antibody thus detects both the full-length as well as the AD-deletion proteins. We observed accumulation of proteins that were of the appropriate size for all transgenic lines (Fig. 1B). These data demonstrate that the full-length and AD-deficient proteins were expressed and specifically detected by the antibody. We quantified signals from three independent transgenic lines for each construct (Fig. 1B, C). As expected, there is variability in expression between individual lines, which can be attributed to position effects of the transgene insertion site. In sum, these analyses demonstrate that all three constructs were successfully expressed in a CP31A-deficient background.

Since CP31A has been shown to support RNA editing at 13 specific sites (Tillich et al., 2009), we investigated these and additional sites in our complementation lines by performing next-generation sequencing of cloned amplified cDNA (Bentolila et al., 2013), testing a total of 16 sites on 10 amplicons. This included sites that were previously shown not to be affected in *cp31a* mutants. Two wt and two *cp31a-1* null mutant plants served as controls. The lowest read coverage observed for any individual site was above 500 reads per site and was found for site *ndhD* 116281 (number refers to the position in the *Arabidopsis* chloroplast genome), while the average coverage for all sites and all genotypes analyzed was 3132 reads per site.

Of the 13 sites previously described to be dependent on CP31A, we confirmed that RNA editing of 11 sites was reduced in *cp31a*-1 mutants (Fig. 2A). Additional defects were found for two previously unreported sites in *ndhB* (94999, 96579), whereas no defect was observed for the previously reported sites, *petL* (65716) and *ndhB* (95225). We speculate that these differences reflect differences in growth conditions: our plants were grown under long-day conditions (16-h light/8-h dark), whereas the previous study used short-day conditions (8-h light/16-h dark; Tillich et al., 2009). Importantly, all our complementation lines, irrespective of the utilized promoter or the presence of the AD showed wt-like editing states at most analyzed sites (Fig. 2A).

**Figure 2:**
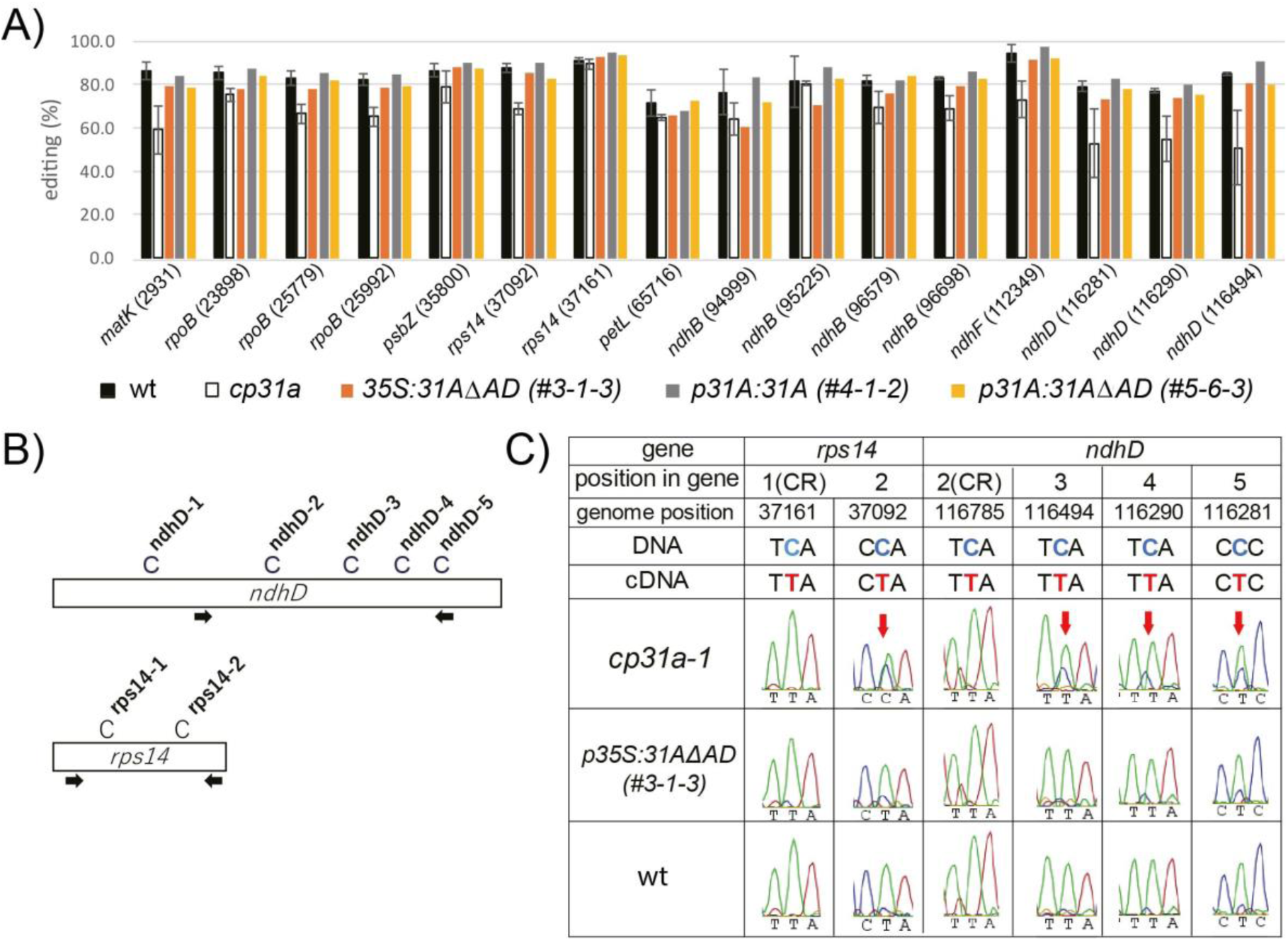
Analysis of RNA editing sites in *cp31a* complementation lines. A) Summary of results from amplicon sequencing of RNA editing sites in complementation lines. Editing sites were amplified by PCR and sequenced on an Illumina platform using barcoded adapters that allowed to eliminate duplicate amplifications. Each individual cDNA sequence is scored for its editing status and the frequency of edited versus non-edited cDNAs is shown. Error bars represent the standard deviation calculated from two replicate experiments with WT and *cp31a* mutants, respectively. Numbers refer to the position of the editing site in the *Arabidopsis thaliana* chloroplast genome. B) Schematic overview of two genes with their editing sites that were assessed by Sanger sequencing of amplified cDNAs. Arrows indicate positions of primers used for cDNA amplification. C) Excerpts from electropherograms showing fluorescence signals for base triplets with the editing site always at the center. Bases marked by red arrows are targets of CP31A. Green traces refer to T signals, blue to C signals and red to A signals. The expected edited (on cDNA) and unedited (on DNA) triplets are shown above. CR = control sites not affected by CP31A according to Tillich et al (2009).

We confirmed this finding by independent amplification and Sanger sequencing of selected RNA-editing sites of the *ndhD* and *rps14* mRNAs. The amplicons we sequenced harbored both, CP31A-dependent and CP31A-independent editing sites (Fig. 2B). Our sequencing confirmed that in *cp31a* mutants, sites *ndhD*-3, 4 and 5, *rps14*-2 show reduced editing efficiency when compared to wt. This defect is complemented in the AD-less CP31A protein (Fig. 2C). The successful complementation demonstrates that the RRM domains are sufficient for the RNA-editing function of CP31A.

### The AD of CP31A is not essential for, but is supportive of the stability of the *ndhF* mRNA

We next analyzed the RNA-stabilizing function of CP31A by focusing on *ndhF*, which is its major target under normal growth conditions (Tillich et al., 2009). Using RNA gel-blot hybridization, we quantified the amount of *ndhF* mRNA in the transgenic lines. The accumulation of this transcript is at least partially restored in all complementing lines relative to the *cp31a* mutant (Fig. 3A, B). Even in a line with very little CP31A accumulation and lacking the AD, a signal for *ndhF* is visible (#3-3-3. Fig. 1, Fig. 3A,B). This indicates that the RRM domains were sufficient to stabilize the *ndhF* mRNA. However, we observed marked differences in the efficiency of *ndhF* stabilization between the full-length and AD-deficient proteins. Lines accumulating large amounts of the AD-less CP31A protein had only little more *ndhF* mRNA than lines expressing much less full-length CP31A protein (compare for example line #3-1-3 with #4-17-1 in Fig. 1B and Fig. 3A). This was visualized by calculating the ratio of *ndhF* mRNA levels to CP31A protein levels (i.e., the stabilization efficiencies; Fig. 3C), which clearly showed that the full-length proteins outperformed the AD-less proteins in stabilizing the *ndhF* mRNA. Together, these results show that the AD supports but is not essential for stabilizing the *ndhF* mRNA and that the RRM domains are capable of performing this task by themselves.

**Figure 3:**
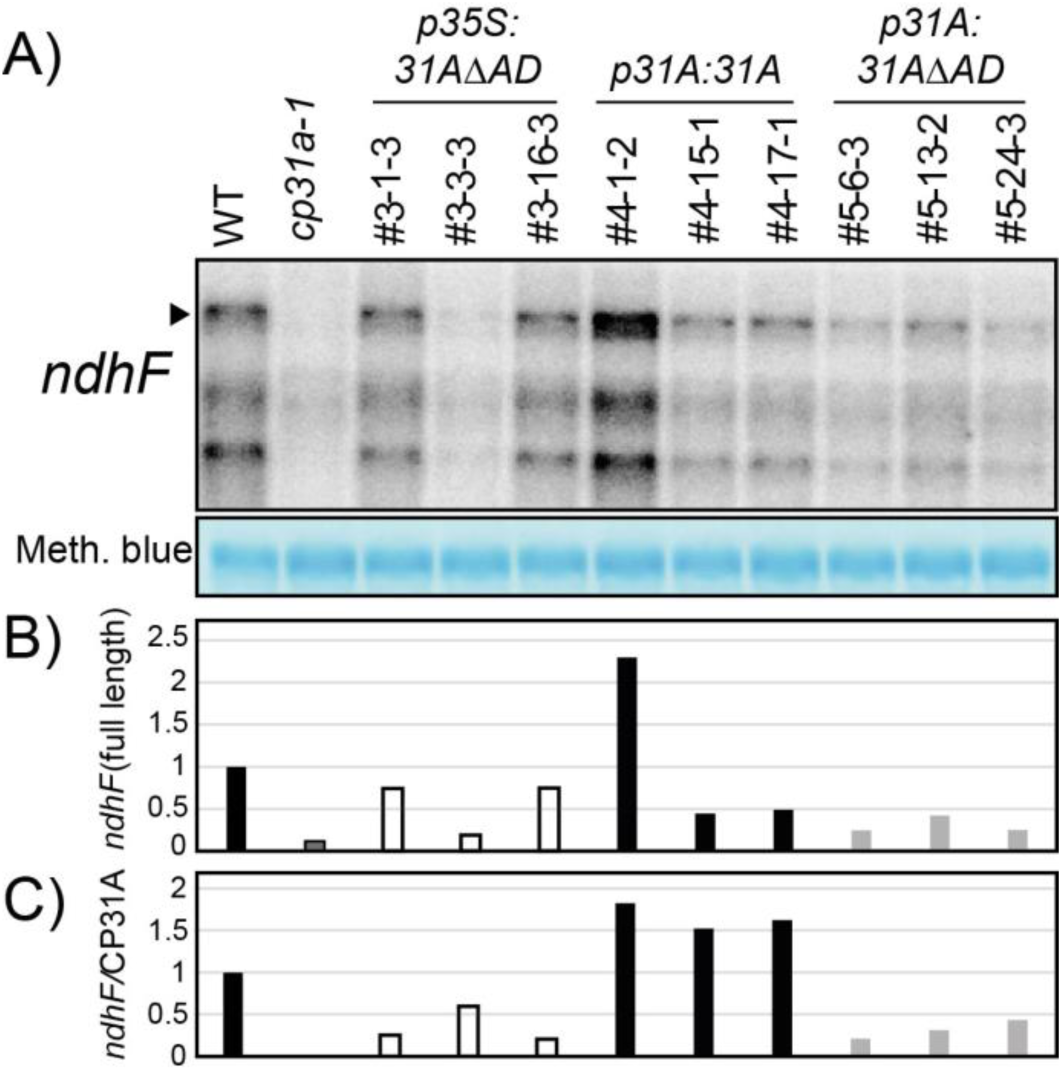
Analysis of the accumulation of the *ndhF* mRNA in *cp31a* complementation lines. A) RNA gel blot analysis of the *ndhF* mRNA in various *cp31a* complementation lines. 4 μg total cellular RNA were separated by denaturing agarose gel electrophoresis, transferred to a nylon membrane and probed for *ndhF*. The transcript labelled with an arrowhead represents the full-length *ndhF* mRNA, whereas smaller signals correspond to degradation products (Kupsch et al. 2012). The nylon membrane was labeled with methylene blue to control for equal loading (excerpt with 23S rRNA shown below blot). B) Quantification of the full-length *ndhF* transcript labeled with an arrowhead in (A). C) Ratio of *ndhF* mRNA levels over CP31A protein levels (from Fig. 1C).

### The AD is required for cold-stress tolerance

To test the impact of the AD on CP31A-mediated cold tolerance, we challenged 2-week-old plants from the transgenic lines with low temperature (8°C) for 5 weeks. Under this treatment, the *cp31a*-null lines are known to display bleaching of the freshly emerging tissue at the center of the leaf rosette (Fig. 4A). Similar losses of pigment were also found in the lines expressing the AD-less CP31A, but not in wt plants or in mutant plants complemented with the full-length protein (Fig. 4A). The pigment deficiency has been shown to be paralleled by a global reduction of the chloroplast-encoded proteins required for photosynthesis in *cp31a* mutants (Kupsch et al., 2012). This reduction of photosynthetic proteins was also observed in plants expressing the AD-less CP31A proteins, while plants expressing full-length CP31A accumulated protein amounts similar to those seen in wt plants (Suppl. Fig. 1). Next, we tested the accumulation of CP31A and *ndhF* in these lines under cold exposure. We separately harvested full leaves, as well as the bleached center part of rosettes in case molecular defects are restricted to the phenotypically conspicuous tissue. However, both tissues showed the same tendency: In line with the findings at normal growth temperatures, the *ndhF*/CP31A ratio was consistently higher for plants expressing full-length CP31A versus plants expressing the AD-less protein (Fig. 4B/C and Suppl. Fig. 2). These experiments show that the AD of CP31A supports the accumulation of chloroplast RNA and is essential for conferring cold resistance to emerging leaf tissue in *Arabidopsis*.

**Figure 4:**
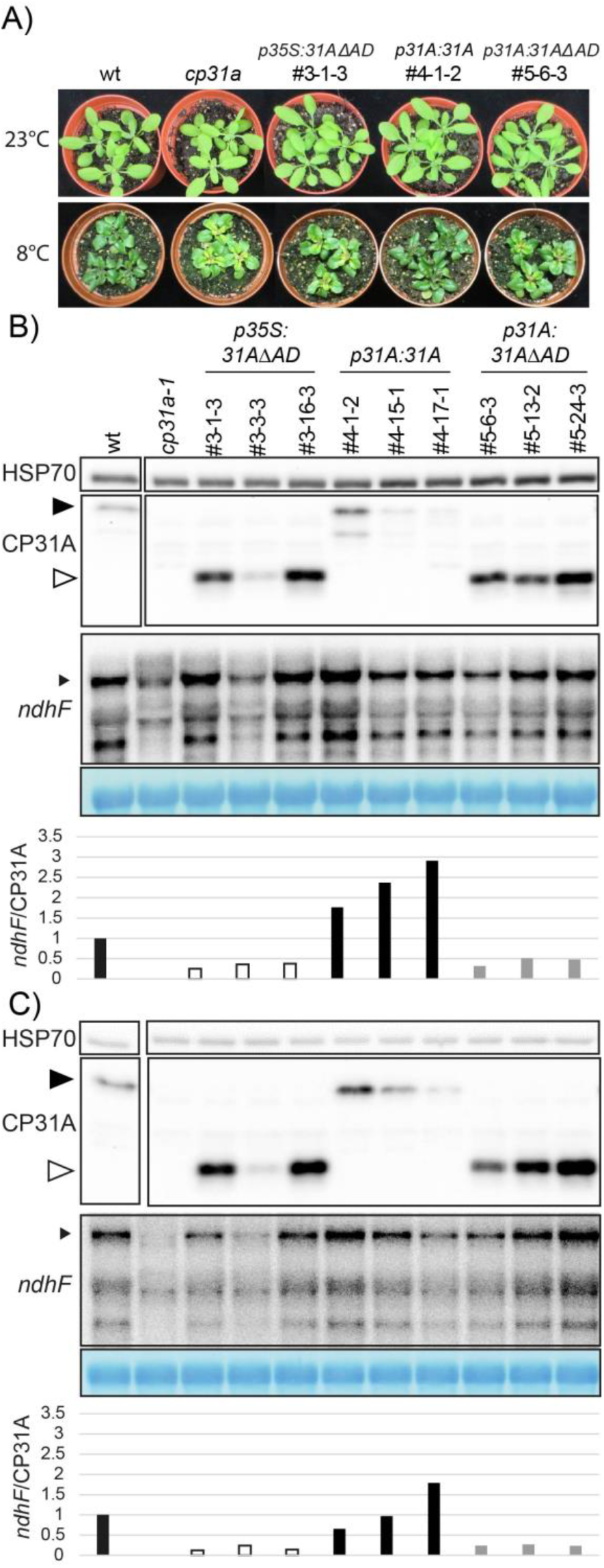
Analysis of *cp31a* complementation lines during cold acclimation. A) Phenotypes of complementation lines grown at 23°C or challenged with 8°C for five weeks. Note the bleached tissue in the center of the rosette in *cp31a* deficient plants and in plants lacking the AD. B) Analysis of CP31A and HSP70 protein accumulation and *ndhF* mRNA accumulation in *cp31a* complementation lines. Here, the entire rosette leaves were used for protein and RNA extraction. Panels from top to bottom: immunoblot analysis of HSP70 proteins as loading control; immunoblot analysis of CP31A proteins; RNA gel blot analysis of *ndhF*; Methylene blue stain of cytosolic rRNA as loading control; Ratio of quantified *ndhF* and CP31A signals. Filled and open arrowheads indicate full-length and AD-less CP31A, respectively, while the small arrowhead denotes the full-length *ndhF* transcript. C) Same analysis as in (B), but with tissue only from the center of the rosette (bleached area in *cp31a* null mutants).

### CP31A associates with multiple mRNAs with a preference for transcripts encoding subunits of the NDH complex, and is required for *ndh* mRNA accumulation at standard growth temperatures

RBPs of various origins and organisms show differential expression under temperature changes, and genetic studies indicate that they are important for temperature-dependent RNA processing (Lin et al., 2010; Malay et al., 2011; Gotic et al., 2016). That said, few studies have examined whether the binding of RBPs to their RNA targets is modified in response to external and internal cues *in vivo*. Since CP31A is required for cold resistance, we tested whether its ability to bind RNA is modified at low temperature and also assayed the importance of the AD for this response. Previous studies using low-resolution techniques could only identify whole transcripts as targets of CP31A (Kupsch et al., 2012). We here used a RIP-Seq approach to obtain better resolution (Suppl. Fig. 3A). For this, wt *Arabidopsis* seedlings and plants expressing the AD-less CP31A protein in a *cp31a* background were grown for 14 days under normal growth conditions and then subjected to formaldehyde cross-linking. For cold tests, seedlings were grown for 13 days at standard growth temperature (21°C), transferred to 4°C for 24 hours, and were then subjected to cross-linking and immunoprecipitation (IP; Kupsch et al., 2012). We used such a short incubation time in the cold to avoid pleiotropic effects that can be expected in the bleached tissue resulting from longer cold challenges. The efficiency of precipitation was comparable between samples grown at normal and low temperatures (Suppl. Fig. 3B). RIP-Seq libraries were prepared in duplicate from input and pellet samples (Suppl. Fig. 3C). We tested the reproducibility of the RIP-Seq experiments by calculating pairwise correlation coefficients across all samples (Suppl. Fig. 3D). We found strong correlations among the input samples (average Pearson coefficient (R): 0.95), and the pellet samples from the IP (average Pearson coefficient (R): 0.97). As expected, input and pellet samples formed separate clusters in our correlation analysis (Suppl. Fig. 3D). This analysis demonstrates high reproducibility between biological replicates of our RIP-Seq assay.

For the analysis of RIP-Seq reads, we followed established procedures originally developed for ChIP-Seq (Bardet et al., 2011; Muino et al., 2011). We started by identifying transcript regions with an enrichment of RNA in pellet samples over input samples. In total, 75 different binding regions were identified as significant in at least one of the four experiment, which are henceforth called binding sites (BSs). The 75 BSs of CP31A are located in 44 transcripts of diverse functionality. Of these 44 transcripts, 31 were also identified by previous RIP-Chip experiments (Suppl. Tab. 1, Suppl. Fig. 4). The difference observed is likely due to methodological dissimilarities. Purified chloroplast stroma was used for the previous RIP-Chip experiments without a cross-link, while here, we used cross-linked frozen total leaf tissue in the RIP-seq. Given these technical differences, it was gratifying to see a similar set of transcripts enriched in the two types of RIP experiments (Suppl. Tab. 1). We next focused on those BSs located within a gene body and thus can be clearly assigned to a specific gene. By contrast, binding sites within intergenic regions cannot be easily assigned to individual genes due to the polycistronic nature of chloroplast transcripts. Assigned BSs are found in genes for the photosynthetic complexes as well as for the gene expression apparatus (Tab. 1). We compared the actual distribution of BSs with the calculated numbers of BSs expected for each functional category (if random binding in the transcriptome is assumed; Tab. 1). This demonstrated that the NDH complex is the only functional category overrepresented among RIP-Seq targets (12 found versus 3 expected; Tab. 1). We also found an unexpected large number of binding sites antisense to coding regions of known genes (Tab. 1). Although antisense RNAs are known to exist in chloroplasts, their levels do not (with a few exceptions) reach those of sense RNAs and their functions (if any) remain unclear (Nakamura et al., 2003; Hotto et al., 2010; Hotto et al., 2011; Sharwood et al., 2011). Consequently, the functional significance of CP31A’s association with antisense RNAs remains unclear as well.

**Table 1:** Summary of target sites of CP31A across all samples analyzed in RIP-Seq experiments

|  | Unique CP31A binding sites<sup>1</sup> |  | Unique CP31A target genes<sup>2</sup> |  | Length of potential target genes (bp)<sup>3</sup> |  | Expected unique binding sites<sup>4</sup> |  |
| --- | --- | --- | --- | --- | --- | --- | --- | --- |
|  | sense | antisense | sense | antisense | sense | antisense | sense | antisense |
| NDH complex | 12 | 0 | 6 | 0 | 13002 | 13002 | 3 | 3 |
| Photosynthesis (except *ndh* genes) | 6 | 3 | 5 | 3 | 23644 | 23644 | 6 | 6 |
| Other metabolism | 4 | 0 | 2 | 0 | 4580 | 4580 | 1 | 1 |
| Ribosome | 0 | 5 | 0 | 5 | 22778 | 22778 | 6 | 6 |
| RNA polymerase | 3 | 1 | 3 | 1 | 11170 | 11170 | 3 | 3 |
| tRNA | 2 | 1 | 2 | 1 | 10196 | 10196 | 2 | 2 |
| Other | 2 | 9 | 2 | 4 | 24177 | 24177 | 6 | 6 |
| All BS | 29 | 19 | 20 | 14 |  |  | 27 | 27 |
| Total BSs in intergenic regions | 27 |  | NA |  | 44931 |  | 21 |  |
| Total BSs | 75 |  | 34 |  |  |  | 75 |  |

<sup>1</sup> this includes binding sites identified in all samples; rRNAs were excluded from calling binding sites
<sup>2</sup> gene annotations according to Araport11
<sup>3</sup> sum of the length of all genes within a category
<sup>4</sup> number of target sites expected in a gene class if the 75 binding sites are randomly distributed in the entire chloroplast genome and if the abundance of transcripts is considered equal between all genes (i.e. depends only on the combined length of all genes within a class)

It is however interesting that at least for the main target category, the *ndh* genes, a preference for sense sites is evident (Tab. 1), suggesting that CP31A is relevant for *ndh* mRNA expression. We therefore analyzed chloroplast RNA levels in *cp31a* versus wt samples at normal temperatures using an oligonucleotide tiling array that represents the entire chloroplast genome of *Arabidopsis thaliana*. We scored only probes whose RNA levels were at least one third lower in the mutant compared to wt. Of all exon probes in the array, 16% represent *ndh* sequences. Importantly, among exon probes whose signals were decreased by at least 0.6-fold in the mutants, 75% contained *ndh* sequences (Fig. 5A, B). No other functional category showed enrichment in this analysis. To confirm these results, we performed RNA gel-blot hybridization experiments for four *ndh* genes that represent the four *ndh* operons in the chloroplast genome (Fig. 5C). We analyzed *cp31a* mutant RNAs alongside RNAs from seedlings with impairment in the expression of *SIGMA FACTOR 4* (*SIG4*), which is specifically required for the transcription of the *ndhF* mRNA (Favory et al., 2005), and *CHLORORESPIRATORY REDUCTION 2* (*CRR2*), which is required for the accumulation of monocistronic *ndhB* transcripts (Hashimoto et al., 2003). As expected, the control mutants showed specific defects for their known target RNAs (Fig. 5C). By contrast, the *ndhK* transcripts accumulated to normal levels in these two control lines, and the *ndhD* mRNA was only slightly decreased (Fig. 5C). In contrast and consistent with our microarray results, the *cp31a* mutants displayed reductions in all four *ndh* transcripts analyzed. The decrease was most pronounced for *ndhF*, strong for *ndhK* and *ndhB*, and somewhat weaker for *ndhD* (Fig. 5C). Collectively, these findings indicate that CP31A stabilizes mRNAs from all four *ndh* operons, suggesting that CP31A regulates *ndh* mRNAs as a group.

**Figure 5:**
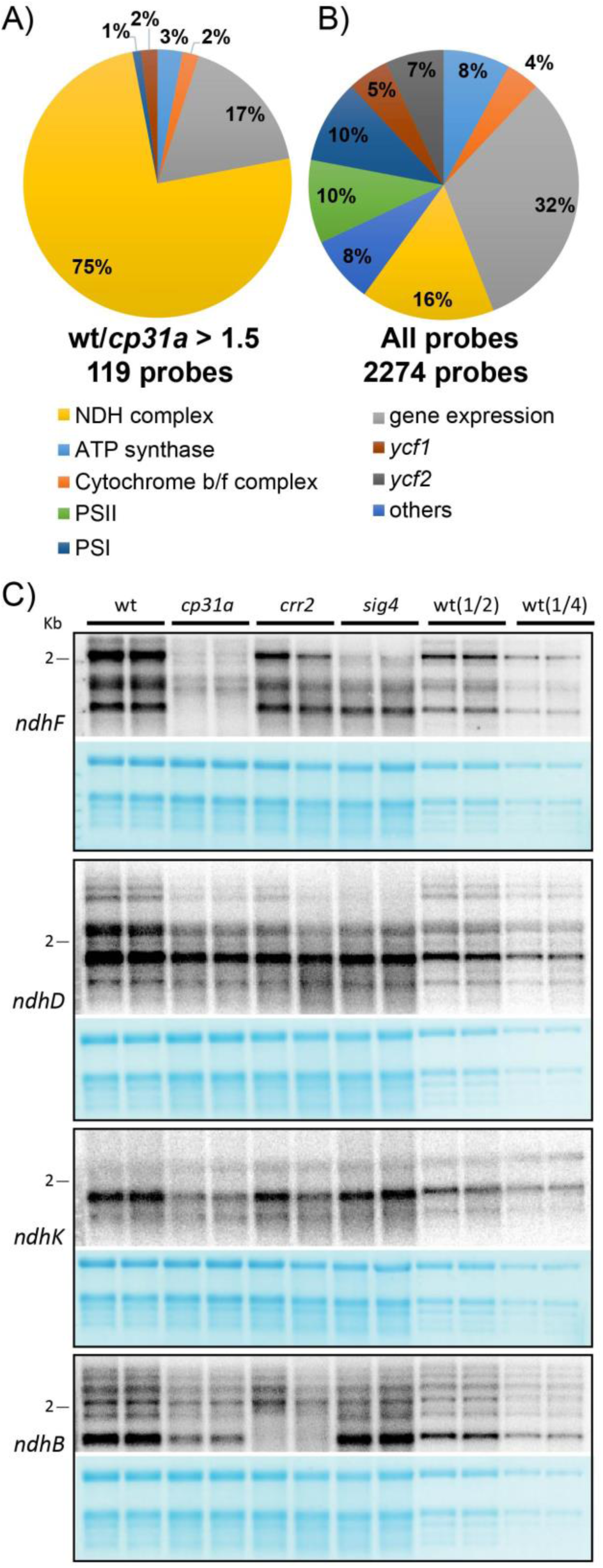
Analysis of RNA accumulation in *cp31a* mutants. A) Summary of microarray analyses of 14 days old wt and *cp31a* mutant seedlings. Relative abundance of exon probes showing at least 1.5 fold stronger signals in wt versus *cp31a* mutants. Three replicate microarray hybridizations were analyzed and assigned to different gene categories. B) Relative distribution of all exon probes on the microarray to different gene categories. C) RNA gel blot analysis of 14 days old Arabidopsis seedlings. 4 μg RNA from from wt, *cp31a* mutants, and two control mutants (*crr2* and *sig4*) together with dilutions of wt samples (1/2 and 1/4) were probed with radiolabeled RNA probes against four different *ndh* genes. The resulting autoradiographs are always shown with the corresponding methylene blue stains of the membranes (below). The 2 kb marker band is shown as a reference.

### *Arabidopsis* CP31A prefers U-rich sequences *in vivo*

Prior *in vitro* experiments showed that a CP31A homolog in tobacco prefers homopolymers of G and U (Li and Sugiura, 1991). We asked whether a sequence preference can be identified in our dataset, i.e. *in vivo*, as well. We used the SSMART motif finder (Munteanu et al., 2018) to query BSs in our RIP-Seq data for common sequences. This uncovered consensus motifs with a clear preference for Us (Fig. 6A; Suppl. Fig. 5). We next compared the frequencies of all possible trinucleotides within significant CP31A BSs against those in all chloroplast coding sequences. Indeed, we found that UUU was the most strongly enriched trinucleotide in this comparison, followed by a number of other U-rich trinucleotides (Fig. 6B). We therefore conclude that CP31A preferentially binds U-rich sequences. As U-rich trinucleotides are the most common trinucleotides in the AT-rich chloroplast genome, our findings suggest that CP31A has evolved to allow a broad RNA target range in the chloroplast transcriptome.

**Figure 6:**
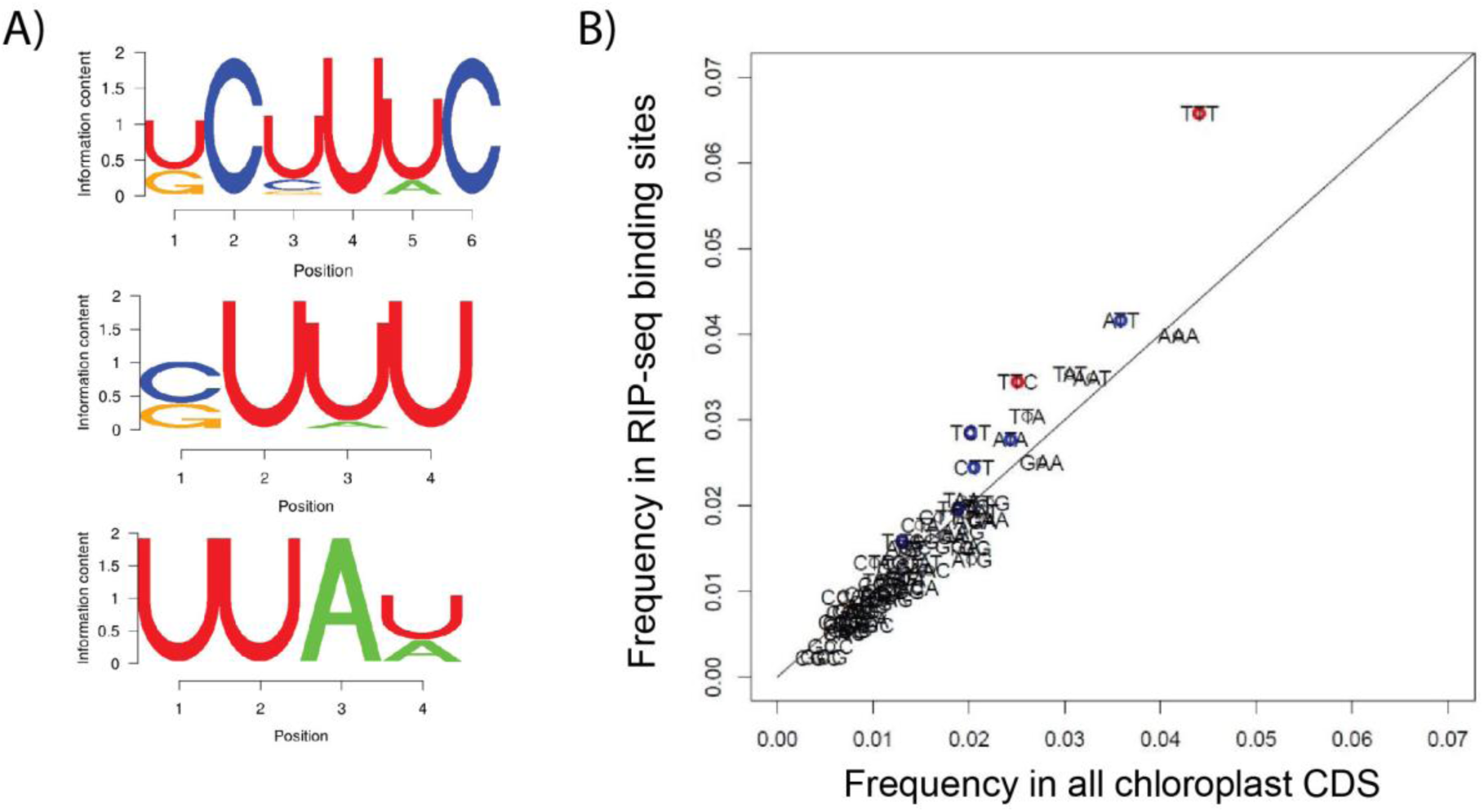
Analysis of sequence elements in CP31A BSs. A) Top scoring consensus motifs identified in CP31A BSs. The SSMART RNA motif finder was run on all significant BSs detected in wt plants grown in cold conditions, as under these conditions, the number of BSs was the highest. The contribution of each BS to the consensus motifs was calculated according to its adjusted p-value - the more significant the BS, the stronger its impact on the consensus. B) Overrepresentation of the UUU(TTT) trinucleotide in CP31A BSs. The scatterplot shows the frequency of all possible trinucleotide sequences in the CP31A BSs regions (y-axis) versus the trinucleotide frequency in all chloroplast coding sequences. Trinucleotides with significant enrichment in RIP-Seq BS regions are marked in red (p<0.01) or blue (p<0.05).

### CP31A shows an increased association with RNA after cold exposure

We hypothesized that CP31A might impact cold resistance in *Arabidopsis* plants through temperature-dependent changes in its affinity for some or all of its target RNAs, which could affect their stability, processing, and/or translation. We therefore compared BSs according to how reliable their enrichment in a given dataset is, based on its FDR value. Using the FDR values, we first focuses on the effect of cold on RNA binding of full-length CP31A and then analyzed the effect of the loss of the AD on binding (next chapter). A majority of BSs were found with high significance in CP31A precipitates in the cold, but were not significantly enriched at higher temperature. This was already visible for some sites in the coverage graphs, as shown in Fig. 7A for binding sites within the *rpl2* mRNAs, or for a site in the intergenic region between the *rps14* and *psbZ* genes. Many BSs show at least some read coverage at both temperatures, but the FDR analysis helped to separate significant from non-significant BSs. The FDR level is displayed for all 75 binding sites as a heat map in Suppl. Fig. 6 and is summarized in Fig. 7B. Most importantly, 30 BSs show significant enrichment with full-length CP31A at 4°C, but not at 21°C (Fig. 7B). Only 9 sites show the opposite behavior, that is, significant enrichment only at 21°C, but not at 4°C. Thus many more sites showed increased versus decreased association with CP31A after cold exposure. Three of the BSs gained in the cold were located in the *ndhF* mRNA, which is known to be a prime target of CP31A (Kupsch et al., 2012). Four more CP31A BSs in *ndh* transcripts were cold-dependent as well (Suppl. Fig. 6). In general, *ndh* mRNAs appeared to be important targets of CP31A under both temperature conditions (Suppl. Fig. 6). Taken together, the results of our temperature-dependent RIP-Seq analysis demonstrate that RNA association of full-length CP31A is higher under cold exposure than at normal temperature, and that *ndh* mRNAs are prime targets under both normal and low temperatures.

**Figure 7:**
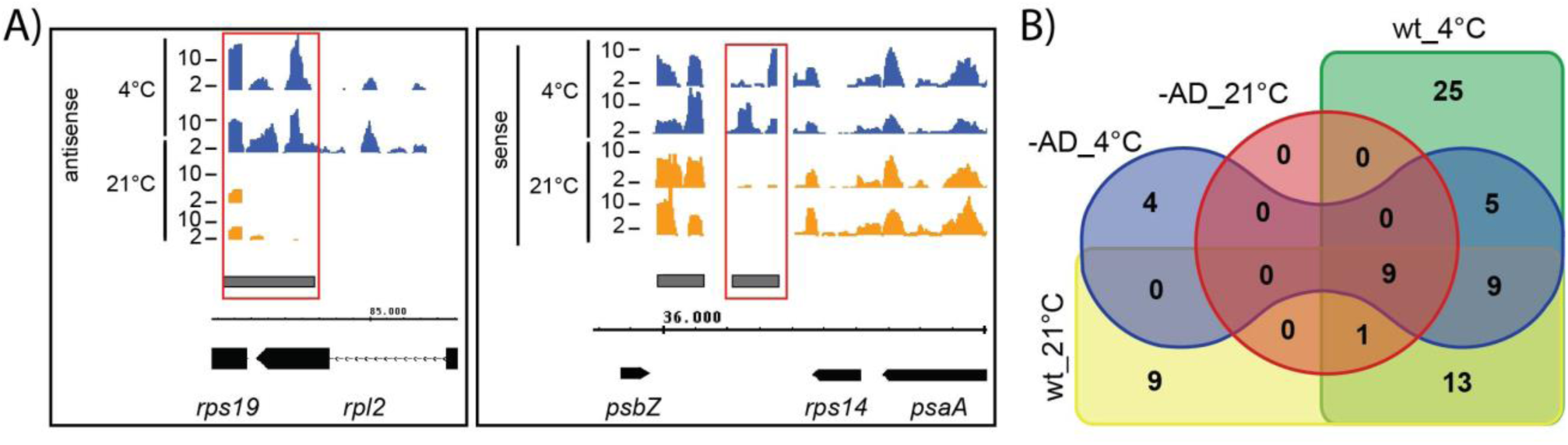
RIP-Seq uncovers cold-induced RNA binding of CP31A. A) Visualization of three examples of temperature-dependent CP31A binding sites. Top track shows the ratio of precipitated versus input reads of RIP-Seq experiments of wt plants grown at 4°C (blue) or 21°C (orange, for two biological replicates at each growth temperature). The track below represents in grey boxes the identified binding sites. The bottom track shows the gene models in black and an intron in *rpl2* as a hatched line. The numbers on the thin black line refer to positions on the *Arabidopsis* chloroplast genome. The red rectangles indicate temperature-dependent CP31A BSs. B) Venn diagram summarizing the occurrence of binding sites in the two genotypes and the two temperature regimes used for growing the plants. For each genotype-temperature combination, only BSs with an FDR < 0.05 were counted.

### The AD of CP31A supports cold-dependent RNA binding

The RRM domain is a highly versatile protein domain that functions in the context of adjacent protein regions. Often, linkers between RRMs or protein domains outside the RRM domains support RNA interactions in addition to the canonical RRM contacts (Lunde et al., 2007). Given this, we used RIP-Seq to test whether the RNA-binding behavior of the AD-less CP31A protein differed from that of the full-length protein. Using the same strategy described for our binding-site analysis with the full-length protein, we scored significant BSs in plants expressing the AD-less variant grown under both normal and cold conditions. We found that the AD-less version of CP31A bound far fewer RNAs than its full-length counterpart (Fig. 7B, Suppl. Fig. 6). Under normal growth temperatures, we identified 41 significant BSs (FDR < 0.05) for the full-length protein and only 10 for the AD-less version (Fig. 7B). Interestingly, the AD-less protein is still responsive to cold: The AD-less protein associated with 18 sites in the cold that are not found to co-precipitate significantly with the protein at normal temperatures (Fig. 7B). On the other hand, AD-less CP31A bound 35 fewer sites than the full-length protein after cold exposure (Fig. 7B). This demonstrates that the AD supports RNA binding under both conditions. Together, our results show (i) that the two RRM domains still react to lowered temperatures in their ability to bind RNA, and (ii) that the AD substantially supports but is not essential for RNA binding.

## Discussion

### CP31A co-regulates *ndh* genes

During the evolution of chloroplasts from cyanobacterial ancestors, operon structures were disrupted and operons were shuffled. Many chloroplast operons therefore include genes with different functions. The *ndh* genes, for example, are separated into four transcriptional units in *Arabidopsis* and are mixed with genes from other functional categories. This lack of conservation of operon structures suggests that transcriptional units are less important than other processes for *ndh* gene regulation in the chloroplast context, giving way to post-transcriptional processes. Translation plays an important role for regulation (Zoschke and Bock, 2018), but it remains unclear, how post-transcriptional co-regulation is achieved prior to translation. We herein demonstrate that CP31A associates with many *ndh* mRNAs under both normal and low temperatures. All analyzed *ndh* mRNAs were reduced in *cp31a* mutants under normal growth conditions, indicating that the CP31A-*ndh* interaction is functionally important. Since the loss of CP31A does not impede transcription (Kupsch et al., 2012), we conclude that the protein is required for the stability of *ndh* mRNAs. An alternative explanation is that the loss of one RNA of the NDH complex has an indirect, hierarchical effect on the other *ndh* RNAs that yields the observed reduction in all *ndh* mRNAs. This phenomenon has been well described for hierarchical protein synthesis cascades in chloroplasts (Choquet et al., 2001), but it has not been previously shown to impact mRNA synthesis or stability. Moreover, the likelihood of this scenario is weakened by our observation that the losses of *ndhF* or *ndhB* in the *sig4* or *crr2* mutants, respectively, were not followed by the loss of all *ndh* mRNAs. Taken together, our data indicate that CP31A combines ndh mRNAs from different genomic loci into a post-transcriptional operon in the chloroplast. Similar post-transcriptional operons, or “RNA regulons”, have been well-described in fruit flies, budding yeast, and mammalian cells (Keene, 2007). In most cases, these RNA operons function in the combined translation and/or stabilization of the participating RNAs (Gerber et al., 2004; Lykke-Andersen and Wagner, 2005; Townley-Tilson et al., 2006). Given the RNA-stabilizing roles of cpRNPs in general and of CP31A in particular, we propose that CP31A adjusts the stability of the *ndh* transcripts as a group.

### The AD contributes to the ability of CP31A to recognize RNA

CP31A has two canonical RRM motifs, which contain the expected aromatic amino acid residues in the RRM’s RNP1 and RNP2 motifs known to be crucial for RNA binding (Ruwe et al., 2011). Here, we show that these RRMs are sufficient to edit chloroplast RNAs and stabilize the *ndhF* mRNA. Still, compared to the wt protein, more AD-less protein is needed to stabilize a similar level of *ndhF* mRNA. Moreover, loss of the AD decreases the ability of CP31A to bind RNA. Thus, the AD domain supports but is not essential for the RNA-binding and RNA-stabilizing functions of CP31A. We do not yet know how the AD achieves its effects at the molecular level. Two non-exclusive possible modes of action are (i) a direct involvement in protein-RNA interactions, and (ii) a role in protein-protein interactions for the recruitment of additional RNA processing factors. Both modes of action were already described in non-plant systems for acidic domains (Lunde et al., 2007). Such functional diversity of an AD has been described for two structural relatives of CP31A, the non-plant RNA binding proteins hnRNP Q and hnRNP R, that also have an N-terminal AD and C-terminal RRM motifs (Geuens et al., 2016). The AD of hnRNP R was shown to support RNA binding and is essential for the role of hnRNP R in the re-initiation of transcription (Fukuda et al., 2013). Similarly, the AD of hnRNP Q forms part of the RNA interaction surface (Hobor et al., 2018) and in addition interacts with the editing factor Apobec (Quaresma et al., 2006). These examples show that it is conceivable that the AD of CP31A contributes to protein-protein interactions (e.g., with other RNA-binding proteins) and also contributes directly to RNA binding. This could serve to increase the affinity for RNA and mediate RNA stabilization and RNA-processing events. A future structural analysis of CP31A bound to RNA would greatly help to understand the mechanistic function of CP31A’s AD.

### Cold-dependent RNA association of CP31A

Many RBPs show differential association with RNA depending on physico-chemical conditions *in vitro*, but relatively little is known about how external and internal cues modulate the target spectrum and target affinity of an RBP *in vivo.* Changed binding to RNA due to external signals has been shown for mammalian RNA binding proteins (Rousseau et al., 2002; Cloutier et al., 2018), but even here, rarely on a genome-wide scale (Benegiamo et al., 2018), and comparable data are missing for plants. Our present results on CP31A show that plant RNA-binding proteins can change their RNA target profiles in response to an external signal, here low temperatures, but it remains unclear how this differential RNA binding is induced. Cold is expected to change RNA structures and CP31A can bind dsDNA *in vitro*, albeit with less affinity than ssDNA and ssRNA (Li and Sugiura, 1991). It is also notable that spinach cpRNPs have been implicated in the establishment of the structured 3’-ends of several mRNAs *in vitro* (Schuster and Gruissem, 1991; Lisitsky et al., 1995; Hayes et al., 1996). These RNA termini require the formation of stem-loop RNA structures, and cpRNP-like proteins have been found to associate with such stem-loops in UV cross-linking experiments (Stern et al., 1989; Chen and Stern, 1991; Danon and Mayfield, 1991; Nickelsen and Link, 1993). Thus, the binding of CP31A to RNAs under cold exposure could reflect the emergence of RNA structures that become more stable at lower temperatures. An *in vivo* analysis of the structure of CP31A target sites under different temperatures could help address the question of why CP31A associates more extensively with RNA in the cold.

What is the functional consequence of the increased RNA association of CP31A under cold exposure? Given that chloroplast RNAs are lost in the bleached tissue of *cp31a* mutants, we propose that an increased interaction with target transcripts could impact the resilience of these transcripts against RNases. An open question however is, which of the many CP31A – transcript interactions in the cold is key to avoid bleaching. Although it is likely that many of these interactions have an incremental effect on the phenotype, we can make some guesses about major contributors. Loss of *ndh* mRNAs, although a prime target of CP31A, is unlikely to be responsible for the bleaching phenotype, since *ndh* genes are downregulated in the cold in *Arabidopsis* (Ivanov et al., 2012), and mutants devoid of the NDH complex do not show bleaching in the cold (Li et al., 2004; Yamori et al., 2011). By contrast, there is a number of other CP31A target RNAs that are known to be essential for chloroplast development and cold-dependent loss of their processing and stabilization could well explain bleaching. This includes mRNAs for subunits of the ATP synthase (atpA, atpB), of the cytochrome *b*_*6*_*f* complex (e.g. *petB, petD*), of the photosystems (e.g. *psaA, psbB*), as well as mRNAs for subunits of the plastid-encoded RNA polymerase (*rpoB, rpoC2*). Loss of the expression of any of these genes leads to defective chloroplasts including pale phenotypes (Hajdukiewicz et al., 1997; Rogalski et al., 2006; Stoppel et al., 2011; Manavski et al., 2015; Chen et al., 2016). Furthermore, many mutants affected in chloroplast translation are more sensitive to cold than wt (Rogalski et al., 2008; Fleischmann et al., 2011; Gu et al., 2014; Wang et al., 2016; Zhang et al., 2016; Paieri et al., 2018; Pulido et al., 2018). This includes mutants of chloroplast RNA binding proteins that affect translation and that show cold-dependent bleaching of the center of the *Arabidopsis* rosette (Wang et al., 2016), which is strikingly similar to the phenotype of *cp31a* mutants in the cold. CP31A binds to several mRNAs coding for ribosomal proteins (e.g. *rps18*), thus being potentially important for the production of the translational machinery in the cold. Taken together, we hypothesize that CP31A becomes important for the expression of a combination of its target mRNAs in the cold; their compromised expression in cold-treated *cp31a* mutants leads to loss of chloroplast development and thus bleaching. Complementation analyses and time-resolved analysis of the accumulation of mRNAs in *cp31a* mutants during cold treatment could reveal, which mRNAs are key targets for CP31A during cold acclimation.

## Methods

### Plant growth

*Arabidopsis thaliana* Columbia-0, *cp31a-1* T-DNA insertion mutants (Tillich et al., 2009) and AD-deletion mutants were grown on soil with a 16-h-light/8-h-dark cycle at 23°C in a CLF growth cabinet at 120 μmol·m^−2^·s^−1^. For cold stress application, temperatures were lowered to 8°C when plants reached an age of two weeks. Cold exposure was then continued for another five weeks to allow the emergence of sufficient new tissue prior to harvest. For RIP-seq experiments plants were grown on a soil/vermiculite 4:1 mixture at 21°C for 14 days (normal conditions). Cold stress was applied after 13 days at 21°C with a 24 h exposure to 4°C.

### Vector construction and production of transgenic plants

A full length *Arabidopsis* CP31A cDNA clone was prepared from total *Arabidopsis* RNA. The genomic region encoding the acidic domain (+241 to +447 bp from the start codon of the open reading frame) was deleted as follows: The 5’ region of *CP31A* (1-240 bp) and the 3’ region of CP31A (448-990 bp) were amplified with the primers 31A_gateway_for and A31-minusADrev and A31-minusAD_for and 31A_gateway_rev, respectively. These 5’ and 3’ regions were joined by restriction-based ligation into a pBluescript vector. The *p35SP:31AΔAD* binary vector for constitutive expression of CP31A was constructed as follows: The *31AΔAD* fragment was amplified with the primer set (SmaI-31AF01; 31A-SmaIR01) and ligated into *pJET1.2* cloning vector (Thermo Scientific), and the *Sma*I digested product was integrated into a *Sma*I site between the *35S* promoter and *Nos* terminator sequences of the binary vector *pGL1* (provided by Dr. Boris Hedtke, HU Berlin), which also contains the bar expression cassette. For producing the *p31AP:31A* vector, the *CP31A* 5’ UTR and promoter region (1,694 bp of the 5’ region of *CP31A*; 12769646 to 12767953 of the *Arabidopsis thaliana* chromosome 4 sequence; Acc. No. CP002687.1) was cloned with the specific primer set (CP31A5’up-1694; CP31A-CDS5’_rev). Then, the *CP31A* cDNA was combined with the promotor by Gibson cloning using the primers (XhoI-31Apro F; Cp31A-CDS5’_rev; CP31A-ADtest_for; 31A-SmaIR01) and cloned into *pJET1.2*, yielding *pXhoI-31AP:31A-SmaI*. The 35S promoter was removed and replaced with multi cloning site including *Xho*I and *Sma*I sites, yielding vector *pGL1-MCS*. XhoI/SmaI digestion products from *pXhoI-31AP:31A-SmaI* were inserted into *Xho*I/*Sma*I sites of *pGL1-MCS*, yielding *p31AP:31A* vector. *p31AP:31AΔAD* was constructed in an analogous way. These three vectors were introduced into the Agrobacterium GV3103 strain independently, and integrated into the knock-out mutant *cp31a-1* (SALK_109613; Tillich et al., 2009) by the floral-dip method. The T_0_ generation was grown on 15 cm diameter plastic plates filled with soil (mixture of horticulture soil and vermiculite). 1/2000 diluted BASTA was sprayed on the plants several times in order to select for BASTA resistant plants. BASTA resistant plants were transferred to plastic pots filled with soil and cultivated at 23°C, 8-h-dark/16-h-light at a light intensity of 120-130 μmol·m^−2^·s^−1^.

### Immunoblot analysis

Total protein (15 μg) were extracted from fully developed leaves grown at normal conditions and immunological analysis were carried out and antibodies were used as previously reported (Kupsch et al., 2012). Immunoblot analysis for the RIP-seq experiments was performed in the same way with protein samples taken from the input, supernatant and pellet fraction of the co-immunoprecipitations.

### RIP-Seq analysis

For RIP-seq experiments *Arabidopsis thaliana* Columbia-0 and AD-deletion mutants under the native promoter (#5-7-1, #5-13-2) were harvested in duplicates after 14 days and flash-frozen in liquid nitrogen. The flash-frozen plant material was grinded under liquid nitrogen to a homogeneous powder. Between 250 to 350 mg plant material was suspended in 3 ml RIP-seq lysis buffer containing formaldehyde (50 mM HEPES-KOH pH 8.0, 200 mM KCl, 5 mM MgCl_2_, 5 mM CaCl_2_, 0.5% Nonidet P-40, 0.5% Sodium deoxycholate, 1x cOmplete^™^, EDTA-free Protease Inhibitor Cocktail (Roche), 100 U RiboLock RNase Inhibitor (ThermoFisher Scientific), 1% formaldehyde) per 1 g plant material. Crosslinking was performed for 10 min under rotation at room temperature and stopped by the addition of 125 mM glycine and a 5 min incubation. The plant extract was centrifuged for 10 min at 20,000xg and 4°C to remove insoluble plant material and flash-frozen in liquid nitrogen.

For the co-immunoprecipitation (CoIP) 8 μl affinity-purified anti-CP31A antibody (Kupsch et al., 2012) was bound to 50 μl Dynabeads ProteinG (Invitrogen) under rotation (15 rpm). The plant extract was thawed and centrifuged for 10 min at 20,000xg and 4°C. 350 μl of the supernatant were diluted 1:1 with CoIP buffer (150 mM NaCl, 20 mM Tris-HCl pH 7.5, 2 mM MgCl_2_, 0.5% Nonidet P-40, 5μg/ml Aprotinin) and incubated with the antibody-coated magnetic beads for 75 min at 15 rpm and 4°C. An aliquot of the antibody-bead solution was taken to serve as the input control. The beads were washed four times in CoIP buffer and resuspended in Proteinase K buffer (100 mM NaCl, 10 mM Tris-HCl pH 7.0, 1 mM EDTA, 0.5% SDS).

The crosslink was reversed with 0.1 mg/ml Proteinase K (ThermoFisher Scientific) at 50°C for 1 h and RNA was extracted from input and pellet fractions using TRIzol and RNA Clean and Concentrator Columns (Zymo Research) according to the manufacturer’s instructions.

Library preparation was performed with the NEBNext^®^ Multiplex Small RNA Library Prep Set for Illumina (New England BioLabs) according to the manufacturer’s instructions with few deviations. Library preparation was performed for half the volume. Additionally, a 5’ adaptor including unique molecular identifiers (UMI) was used (5′-rGrUrUrCrArGrArGrUrUrCrUrArCrArGrUrCrCrGrArCrGrArUrCGATCNNNNNNNN-3’). PCR amplification was performed using the KAPA HiFi HotStart ReadyMix with the cycling protocol for library amplification for Illumina platforms and an annealing temperature of 62°C. The PCR amplified cDNA construct was purified using the GeneJET PCR Purification Kit (ThermoFisher Scientific) according to the manufacturer’s instructions and then separated on 6% polyacrylamide gel. Library fragments between 160bp and 190bp were extracted from the gel according to the NEBNext^®^ protocol and subjected to Illumina sequencing. Read numbers and mapping results are summarized for all libraries in Supple. Table 3.

For the bioinformatic identification of the CP31A binding events the following steps were performed:

1. Mapping of the Illumina reads. First, reads were split depending on its barcode using *fastx_barcode_splitter* (version 11sep2008) to identify to which sample they correspond, next the UMI barcode was extracted for each read using *umi_tools*, later, reads were trimmed using Trimmomatic version 0.36 with the next parameters: *LEADING:3 TRAILING:3 SLIDINGWINDOW:4:15 MINLEN:10* in order to eliminate Illumina adapters and bad quality sequence regions. The obtained trimmed reads were aligned to the *Arabidopsis thaliana* genomes (Araport11) using STAR version 2.5.2a with parameter: *-- outFilterMultimapNmax 2 --outSAMtype BAM SortedByCoordinate --alignIntronMax 3000 -- outFilterIntronMotifs RemoveNoncanonical --outSAMstrandField intronMotif --outSAMattrIHstart 0*. At this point, duplicated reads were removed from the BAM file using *umi_tools.* Only reads that mapped to the chloroplast genome with a length bigger than 35 bp were used for downstream analysis.
2. Identification of candidate BSs in each condition studied. Reads mapped in rRNA were eliminated. Remaining reads were used to identify BSs using CSAR (version 1.31.0; parameters: *b=10, w=70, nper=100, test=“Ratio”*) for each strand and sample/replicate independently. Only those regions that showed a significant enrichment in mapped reads when comparing IP versus corresponding input samples at a false discovery rate (FDR) < 0.05 were kept (Muino et al., 2011), which led to the discovery of 371 candidate BSs across all samples. In order to obtain a common set of candidate BSs for all samples, the list of BSs for each sample/replicate were merged using the software *mergeBed* from the package *bedtools* (v2.26.0). This resulted in 75 reference BSs.
3. Quantification of BSs. The program *featurecounts* (v1.6.0; parameters: *-s 1 -M*) was used to count the number of reads strand-specifically mapping to the common set of candidate BSs obtained from the previous step. DESeq2 (v1.14.1) was used with defaults parameters except *fitType=“local”* to obtain normalized number of mapped read in each sample, next it was used to calculate enrichment and significance of number of the candidate BSs comparing IP vs control (two biological replicates). The normalized number of reads were used to calculate Pearson correlation coefficients.

### SSMART analysis of CP31A RIP-seq data

SSMART analysis was carried out as described (Munteanu et al., 2018). We used all significant BSs identified in wild type plants grown in cold conditions as input, as in this sequence set most binding events were detected. The input sequences of the BSs were scored using their adjusted p-value according to the actual binding site analysis of the RIP-seq data.

### RNA extraction and editing analysis

Total RNA was extracted from fully developed leafs (0.1 g) powdered in liquid nitrogen using Trizol (Thermo Fisher) according to the manufacturer’s protocol. DNA was removed from RNA samples by three consecutive DNase I treatments and Phenol/Chloroform/Isoamyl alcohol extractions. DNA removal was checked by PCR with the chloroplast-specific *rps14* primer set. cDNA synthesis was performed with 2 μg RNA using SuperScript III first strand synthesis system (Invitrogen) according to the manufacturer’s protocol. A one-to-ten dilution of cDNA was used as a template for amplifying ten cDNAs encompassing 16 editing sites (**Fig.** 2A) on nine plastid genes using primer sets previously reported (Tillich et al., 2009). RT-PCR products were purified with MinElute PCR Purification Kit (Qiagen). Bulk RT-PCR products were cloned into the barcoded cloning vector and sequenced using the Illumina MiSeq machine. Sequenced data were sorted according to barcoding of vectors and analyzed with the CLC Genomics Workbench (CLC bio). Selected RT-PCR products were directly analyzed by Sanger Sequencing and analyzed using the Geneious software.

### RNA gel blot analysis

Total RNA (4 μg) was fractioned on 1.2% agarose gels containing 1.2% formaldehyde, blotted and hybridized with radiolabeled RNA probes produced by T7 *in vitro* transcription from PCR products generated with primer combinations described in Suppl. Tab. 2.

## Acknowledgement

We wish to acknowledge expert technical support by Irina Passow. This work was supported by grants of the Deutsche Forschungsgemeinschaft to CSL (A02 of TRR175), DL (C05 of TRR175) and TR (FOR 2092). Support of MKL by the IRI Life Sciences and the “Frauenförderung des Instituts für Biologie” of the Humboldt University is gratefully acknowledged. AO was supported by JSPS Overseas Research Fellowships. We thank Deserah Strand and Hannes Ruwe for critical discussion of the data.

AO, MKL, and TR performed the research and analyzed the data. JMM performed the computational analysis of the RNA seq data. BL performed the analysis of the CP31A BS consensus. DL, UO and CSL designed the research and analyzed the data. CSL wrote the paper with contributions of all authors to the final version.

## References

Bardet AF, He Q, Zeitlinger J, Stark A (2011) A computational pipeline for comparative ChIP-seq analyses. Nat Protoc 7: 45–61

Barkan A (2011) Expression of plastid genes: organelle-specific elaborations on a prokaryotic scaffold. Plant Physiol 155: 1520–1532

Barkan A, Small I (2014) Pentatricopeptide repeat proteins in plants. Annu Rev Plant Biol 65: 415–442

Benegiamo G, Mure LS, Erikson G, Le HD, Moriggi E, Brown SA, Panda S (2018) The RNA-Binding Protein NONO Coordinates Hepatic Adaptation to Feeding. Cell Metab 27: 404–418 e407

Bentolila S, Oh J, Hanson MR, Bukowski R (2013) Comprehensive high-resolution analysis of the role of an Arabidopsis gene family in RNA editing. PLoS Genet 9: e1003584

Biehl A, Richly E, Noutsos C, Salamini F, Leister D (2005) Analysis of 101 nuclear transcriptomes reveals 23 distinct regulons and their relationship to metabolism, chromosomal gene distribution and co-ordination of nuclear and plastid gene expression. Gene 344: 33–41

Bollenbach TJ, Schuster G, Portnoy V, Stern D (2007) Processing, degradation, and polyadenylation of chloroplast transcripts. In R Bock, ed, Cell and Molecular Biology of Plastids, Vol 19. Springer, Berlin, Heidelberg, pp 175–211

Castandet B, Hotto AM, Strickler SR, Stern DB (2016) ChloroSeq, an Optimized Chloroplast RNA-Seq Bioinformatic Pipeline, Reveals Remodeling of the Organellar Transcriptome Under Heat Stress. G3 (Bethesda) 6: 2817–2827

Chen F, Dong G, Wu L, Wang F, Yang X, Ma X, Wang H, Wu J, Zhang Y, Wang H, Qian Q, Yu Y (2016) A Nucleus-Encoded Chloroplast Protein YL1 Is Involved in Chloroplast Development and Efficient Biogenesis of Chloroplast ATP Synthase in Rice. Sci Rep 6: 32295

Chen HC, Stern DB (1991) Specific ribonuclease activities in spinach chloroplasts promote mRNA maturation and degradation. J Biol Chem 266: 24205–24211

Cho WK, Geimer S, Meurer J (2009) Cluster analysis and comparison of various chloroplast transcriptomes and genes in Arabidopsis thaliana. DNA Res 16: 31–44

Choquet Y, Wostrikoff K, Rimbault B, Zito F, Girard-Bascou J, Drapier D, Wollman FA (2001) Assembly-controlled regulation of chloroplast gene translation. Biochem Soc Trans 29: 421–426

Cloutier A, Shkreta L, Toutant J, Durand M, Thibault P, Chabot B (2018) hnRNP A1/A2 and Sam68 collaborate with SRSF10 to control the alternative splicing response to oxaliplatin-mediated DNA damage. Sci Rep 8: 2206

Danon A, Mayfield APY (1991) Light regulated translational activators: identification of chloroplast gene specific mRNA binding proteins. EMBO J. 10: 3993–4001

Daros JA, Flores R (2002) A chloroplast protein binds a viroid RNA in vivo and facilitates its hammerhead-mediated self-cleavage. EMBO J 21: 749–759

Deng X-W, Gruissem W (1987) Control of plastid gene expression during development: the limited role of transcriptional regulation. Cell 49: 379–387

Deng XW, Tonkyn JC, Peter GF, Thornber JP, Gruissem W (1989) Post-transcriptional control of plastid mRNA accumulation during adaptation of chloroplasts to different light quality environments. Plant Cell 1: 645–654

Eberhard S, Drapier D, Wollman F (2002) Searching limiting steps in the expression of chloroplast-encoded proteins: relations between gene copy number, transcription, transcript abundance and translation rate in the chloroplast of *Chlamydomonas reinhardtii*. Plant J. 31: 149–160

Favory JJ, Kobayshi M, Tanaka K, Peltier G, Kreis M, Valay JG, Lerbs-Mache S (2005) Specific function of a plastid sigma factor for ndhF gene transcription. Nucleic Acids Res 33: 5991–5999

Fleischmann TT, Scharff LB, Alkatib S, Hasdorf S, Schottler MA, Bock R (2011) Nonessential plastid-encoded ribosomal proteins in tobacco: a developmental role for plastid translation and implications for reductive genome evolution. Plant Cell 23: 3137–3155

Fukuda A, Shimada M, Nakadai T, Nishimura K, Hisatake K (2013) Heterogeneous nuclear ribonucleoprotein R cooperates with mediator to facilitate transcription reinitiation on the c-Fos gene. PLoS One 8: e72496

Gerber AP, Herschlag D, Brown PO (2004) Extensive association of functionally and cytotopically related mRNAs with Puf family RNA-binding proteins in yeast. PLoS Biol 2: E79

Germain A, Hotto AM, Barkan A, Stern DB (2013) RNA processing and decay in plastids. Wiley Interdiscip Rev RNA 4: 295–316

Germain A, Kim SH, Gutierrez R, Stern DB (2012) Ribonuclease II preserves chloroplast RNA homeostasis by increasing mRNA decay rates, and cooperates with polynucleotide phosphorylase in 3’ end maturation. Plant J 72: 960–971

Geuens T, Bouhy D, Timmerman V (2016) The hnRNP family: insights into their role in health and disease. Hum Genet 135: 851–867

Gotic I, Omidi S, Fleury-Olela F, Molina N, Naef F, Schibler U (2016) Temperature regulates splicing efficiency of the cold-inducible RNA-binding protein gene Cirbp. Genes Dev 30: 2005–2017

Grimmer J, Rodiger A, Hoehenwarter W, Helm S, Baginsky S (2014) The RNA-binding protein RNP29 is an unusual Toc159 transport substrate. Front Plant Sci 5: 258

Gu L, Xu T, Lee K, Lee K, Kang H (2014) A chloroplast-localized DEAD-box RNA helicase AtRH3 is essential for intron splicing and plays an important role in the growth and stress response in Arabidopsis thaliana. Plant Physiol Biochem 82: 309–318

Hajdukiewicz PT, Allison LA, Maliga P (1997) The two RNA polymerases encoded by the nuclear and the plastid compartments transcribe distinct groups of genes in tobacco plastids. EMBO J 16: 4041–4048

Hashimoto M, Endo T, Peltier G, Tasaka M, Shikanai T (2003) A nucleus-encoded factor, CRR2, is essential for the expression of chloroplast *ndhB* in Arabidopsis. Plant J 36: 541–549

Hayes R, Kudla J, Schuster G, Gabay L, Maliga P, Gruissem W (1996) Chloroplast mRNA 3’-end processing by a high molecular weight protein complex is regulated by nuclear encoded RNA binding proteins. EMBO J 15: 1132–1141

Hirose T, Sugiura M (2001) Involvement of a site-specific *trans*-acting factor and a common RNA-binding protein in the editing of chloroplast mRNAs: development of a chloroplast in vitro RNA editing system. EMBO J 20: 1144–1152

Hobor F, Dallmann A, Ball NJ, Cicchini C, Battistelli C, Ogrodowicz RW, Christodoulou E, Martin SR, Castello A, Tripodi M, Taylor IA, Ramos A (2018) A cryptic RNA-binding domain mediates Syncrip recognition and exosomal partitioning of miRNA targets. Nat Commun 9: 831

Hotto AM, Huston ZE, Stern DB (2010) Overexpression of a natural chloroplast-encoded antisense RNA in tobacco destabilizes 5S rRNA and retards plant growth. BMC Plant Biol 10: 213

Hotto AM, Schmitz RJ, Fei Z, Ecker JR, Stern DB (2011) Unexpected Diversity of Chloroplast Noncoding RNAs as Revealed by Deep Sequencing of the Arabidopsis Transcriptome. G3 (Bethesda) 1: 559–570

Ivanov AG, Rosso D, Savitch LV, Stachula P, Rosembert M, Oquist G, Hurry V, Huner NP (2012) Implications of alternative electron sinks in increased resistance of PSII and PSI photochemistry to high light stress in cold-acclimated Arabidopsis thaliana. Photosynth Res 113: 191–206

Keene JD (2007) RNA regulons: coordination of post-transcriptional events. Nat Rev Genet 8: 533–543

Klaff P, Gruissem W (1991) Changes in chloroplast mRNA stability during leaf development. Plant Cell 3: 517–529

Klein RR (1991) Regulation of light-induced chloroplast transcription and translation in eight-day-old dark-grown barley seedlings. Plant Physiol. 97: 335–342

Kupsch C, Ruwe H, Gusewski S, Tillich M, Small I, Schmitz-Linneweber C (2012) Arabidopsis Chloroplast RNA Binding Proteins CP31A and CP29A Associate with Large Transcript Pools and Confer Cold Stress Tolerance by Influencing Multiple Chloroplast RNA Processing Steps. Plant Cell 10: 4266–4280

Li XG, Duan W, Meng QW, Zou Q, Zhao SJ (2004) The function of chloroplastic NAD(P)H dehydrogenase in tobacco during chilling stress under low irradiance. Plant Cell Physiol 45: 103–108

Li YQ, Sugiura M (1990) Three distinct ribonucleoproteins from tobacco chloroplasts: each contains a unique amino terminal acidic domain and two ribonucleoprotein consensus motifs. EMBO J 9: 3059–3066

Li YQ, Sugiura M (1991) Nucleic acid-binding specificities of tobacco chloroplast ribonucleoproteins. Nucleic Acids Res 19: 2893–2896

Lin BC, Defenbaugh DA, Casey JL (2010) Multimerization of hepatitis delta antigen is a critical determinant of RNA binding specificity. J Virol 84: 1406–1413

Lisitsky I, Liveanu V, Schuster G (1995) RNA-Binding Characteristics of a Ribonucleoprotein from Spinach Chloroplast. Plant Physiol. 107: 933–941

Lisitsky I, Schuster G (1995) Phosphorylation of a chloroplast RNA-binding protein changes its affinity to RNA. Nucl. Acids Res. 23: 2506–2511

Loza-Tavera H, Vargas-Suarez M, Diaz-Mireles E, Torres-Marquez ME, Gonzalez de la Vara LE, Moreno-Sanchez R, Gruissem W (2006) Phosphorylation of the spinach chloroplast 24 kDa RNA-binding protein (24RNP) increases its binding to petD and psbA 3’ untranslated regions. Biochimie 88: 1217–1228

Lunde BM, Moore C, Varani G (2007) RNA-binding proteins: modular design for efficient function. Nat Rev Mol Cell Biol 8: 479–490

Lykke-Andersen J, Wagner E (2005) Recruitment and activation of mRNA decay enzymes by two ARE-mediated decay activation domains in the proteins TTP and BRF-1. Genes Dev 19: 351–361

Malay AD, Watanabe M, Heddle JG, Tame JR (2011) Crystal structure of unliganded TRAP: implications for dynamic allostery. Biochem J 434: 427–434

Manavski N, Schmid LM, Meurer J (2018) RNA-stabilization factors in chloroplasts of vascular plants. Essays Biochem 62: 51–64

Manavski N, Torabi S, Lezhneva L, Arif MA, Frank W, Meurer J (2015) HIGH CHLOROPHYLL FLUORESCENCE145 Binds to and Stabilizes the psaA 5’ UTR via a Newly Defined Repeat Motif in Embryophyta. Plant Cell 27: 2600–2615

Mentzen WI, Wurtele ES (2008) Regulon organization of Arabidopsis. BMC Plant Biol 8: 99

Muino JM, Kaufmann K, van Ham RC, Angenent GC, Krajewski P (2011) ChIP-seq Analysis in R (CSAR): An R package for the statistical detection of protein-bound genomic regions. Plant Methods 7: 11

Munteanu A, Mukherjee N, Ohler U (2018) SSMART: sequence-structure motif identification for RNA-binding proteins. Bioinformatics 34: 3990–3998

Nakamura T, Furuhashi Y, Hasegawa K, Hashimoto H, Watanabe K, Obokata J, Sugita M, Sugiura M (2003) Array-based analysis on tobacco plastid transcripts: preparation of a genomic microarray containing all genes and all intergenic regions. Plant Cell Physiol 44: 861–867

Nakamura T, Ohta M, Sugiura M, Sugita M (1999) Chloroplast ribonucleoproteins are associated with both mRNAs and intron-containing precursor tRNAs. FEBS Lett 460: 437–441

Nakamura T, Ohta M, Sugiura M, Sugita M (2001) Chloroplast ribonucleoproteins function as a stabilizing factor of ribosome-free mRNAs in the stroma. J Biol Chem 276: 147–152

Nickelsen J, Link G (1993) The 54 kDa RNA-binding protein from mustard chloroplasts mediates endonucleolytic transcript 3’ end formation in vitro. Plant J 3: 537–544

Paieri F, Tadini L, Manavski N, Kleine T, Ferrari R, Morandini P, Pesaresi P, Meurer J, Leister D (2018) The DEAD-box RNA Helicase RH50 Is a 23S-4.5S rRNA Maturation Factor that Functionally Overlaps with the Plastid Signaling Factor GUN1. Plant Physiol 176: 634–648

Pfannschmidt T (2003) Chloroplast redox signals: how photosynthesis controls its own genes. Trends Plant Sci 8: 33–41

Pulido P, Zagari N, Manavski N, Gawronski P, Matthes A, Scharff LB, Meurer J, Leister D (2018) CHLOROPLAST RIBOSOME ASSOCIATED Supports Translation under Stress and Interacts with the Ribosomal 30S Subunit. Plant Physiol 177: 1539–1554

Quaresma AJ, Oyama S, Jr., Barbosa JA, Kobarg J (2006) The acidic domain of hnRNPQ (NSAP1) has structural similarity to Barstar and binds to Apobec1. Biochem Biophys Res Commun 350: 288–297

Reiland S, Messerli G, Baerenfaller K, Gerrits B, Endler A, Grossmann J, Gruissem W, Baginsky S (2009) Large-scale Arabidopsis phosphoproteome profiling reveals novel chloroplast kinase substrates and phosphorylation networks. Plant Physiol 150: 889–903

Rogalski M, Ruf S, Bock R (2006) Tobacco plastid ribosomal protein S18 is essential for cell survival. Nucleic Acids Res 34: 4537–4545

Rogalski M, Schottler MA, Thiele W, Schulze WX, Bock R (2008) Rpl33, a nonessential plastid-encoded ribosomal protein in tobacco, is required under cold stress conditions. Plant Cell 20: 2221–2237

Rousseau S, Morrice N, Peggie M, Campbell DG, Gaestel M, Cohen P (2002) Inhibition of SAPK2a/p38 prevents hnRNP A0 phosphorylation by MAPKAP-K2 and its interaction with cytokine mRNAs. EMBO J 21: 6505–6514

Ruwe H, Kupsch C, Teubner M, Schmitz-Linneweber C (2011) The RNA-recognition motif in chloroplasts. J Plant Physiol 168: 1361–1371

Schonberg A, Bergner E, Helm S, Agne B, Dunschede B, Schunemann D, Schutkowski M, Baginsky S (2014) The peptide microarray “ChloroPhos1.0” identifies new phosphorylation targets of plastid casein kinase II (pCKII) in Arabidopsis thaliana. PLoS One 9: e108344

Schuster G, Gruissem W (1991) Chloroplast mRNA 3’ end processing requires a nuclear-encoded RNA-binding protein. EMBO J. 10: 1493–1502

Schuster G, Lisitsky I, Klaff P (1999) Polyadenylation and degradation of mRNA in the chloroplast. Plant Physiol 120: 937–944

Selinger DW, Saxena RM, Cheung KJ, Church GM, Rosenow C (2003) Global RNA half-life analysis in Escherichia coli reveals positional patterns of transcript degradation. Genome Res 13: 216–223

Sharwood RE, Hotto AM, Bollenbach TJ, Stern DB (2011) Overaccumulation of the chloroplast antisense RNA AS5 is correlated with decreased abundance of 5S rRNA in vivo and inefficient 5S rRNA maturation in vitro. RNA 17: 230–243

Stern DB, Jones H, Gruissem W (1989) Function of plastid mRNA 3’inverted repeats. RNA stabilization and gene specific protein binding. J. Biol. Chem. 264: 18742–18750

Stoppel R, Lezhneva L, Schwenkert S, Torabi S, Felder S, Meierhoff K, Westhoff P, Meurer J (2011) Recruitment of a ribosomal release factor for light- and stress-dependent regulation of petB transcript stability in Arabidopsis chloroplasts. Plant Cell 23: 2680–2695

Teubner M, Fuss J, Kuhn K, Krause K, Schmitz-Linneweber C (2017) The RNA recognition motif protein CP33A is a global ligand of chloroplast mRNAs and is essential for plastid biogenesis and plant development. Plant J 89: 472–485

Tillich M, Hardel SL, Kupsch C, Armbruster U, Delannoy E, Gualberto JM, Lehwark P, Leister D, Small ID, Schmitz-Linneweber C (2009) Chloroplast ribonucleoprotein CP31A is required for editing and stability of specific chloroplast mRNAs. Proc Natl Acad Sci U S A 106: 6002–6007

Townley-Tilson WH, Pendergrass SA, Marzluff WF, Whitfield ML (2006) Genome-wide analysis of mRNAs bound to the histone stem-loop binding protein. RNA 12: 1853–1867

Tsunoyama Y, Ishizaki Y, Morikawa K, Kobori M, Nakahira Y, Takeba G, Toyoshima Y, Shiina T (2004) Blue light-induced transcription of plastid-encoded psbD gene is mediated by a nuclear-encoded transcription initiation factor, AtSig5. Proc Natl Acad Sci U S A 101: 3304–3309

Udy DB, Belcher S, Williams-Carrier R, Gualberto JM, Barkan A (2012) Effects of reduced chloroplast gene copy number on chloroplast gene expression in maize. Plant Physiol 160: 1420–1431

Wang BC, Wang HX, Feng JX, Meng DZ, Qu LJ, Zhu YX (2006) Post-translational modifications, but not transcriptional regulation, of major chloroplast RNA-binding proteins are related to Arabidopsis seedling development. Proteomics 6: 2555–2563

Wang S, Bai G, Wang S, Yang L, Yang F, Wang Y, Zhu JK, Hua J (2016) Chloroplast RNA-Binding Protein RBD1 Promotes Chilling Tolerance through 23S rRNA Processing in Arabidopsis. PLoS Genet 12: e1006027

Yamori W, Sakata N, Suzuki Y, Shikanai T, Makino A (2011) Cyclic electron flow around photosystem I via chloroplast NAD(P)H dehydrogenase (NDH) complex performs a significant physiological role during photosynthesis and plant growth at low temperature in rice. Plant J 68: 966–976

Ye L, Sugiura M (1992) Domains required for nucleic acid binding activities in chloroplast ribonucleoproteins. Nucleic Acids Res 20: 6275–6279

Zhang J, Yuan H, Yang Y, Fish T, Lyi SM, Thannhauser TW, Zhang L, Li L (2016) Plastid ribosomal protein S5 is involved in photosynthesis, plant development, and cold stress tolerance in Arabidopsis. J Exp Bot 67: 2731–2744

Zoschke R, Bock R (2018) Chloroplast Translation: Structural and Functional Organization, Operational Control, and Regulation. Plant Cell 30: 745–770

