## Supplemental Figures for "Chloroplast cold-resistance is mediated by the acidic domain of the RNA binding protein CP31A"

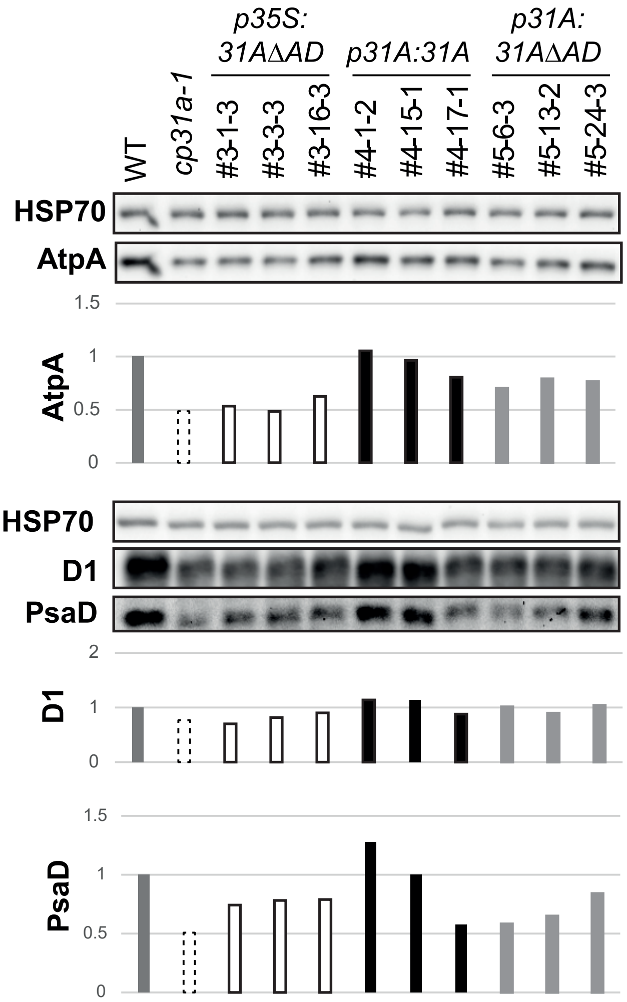


Supplemental Figure 1: Analysis of photosynthetic protein accumulation in *cp31a* complementation lines after cold treatment using total leaf protein preparations. Top panel: immunoblot analysis of the chloroplast ATPA protein and, as a loading control, of the HSP70 protein. Quantification of chemoluminescence signals of D1 relative to HSP70 is shown below. Bottom panels: independent immunoblot analysis of D1 and PsaD, again in relation to the HSP70 signal. All three proteins were detected on the same blot. Quantifications are shown below.

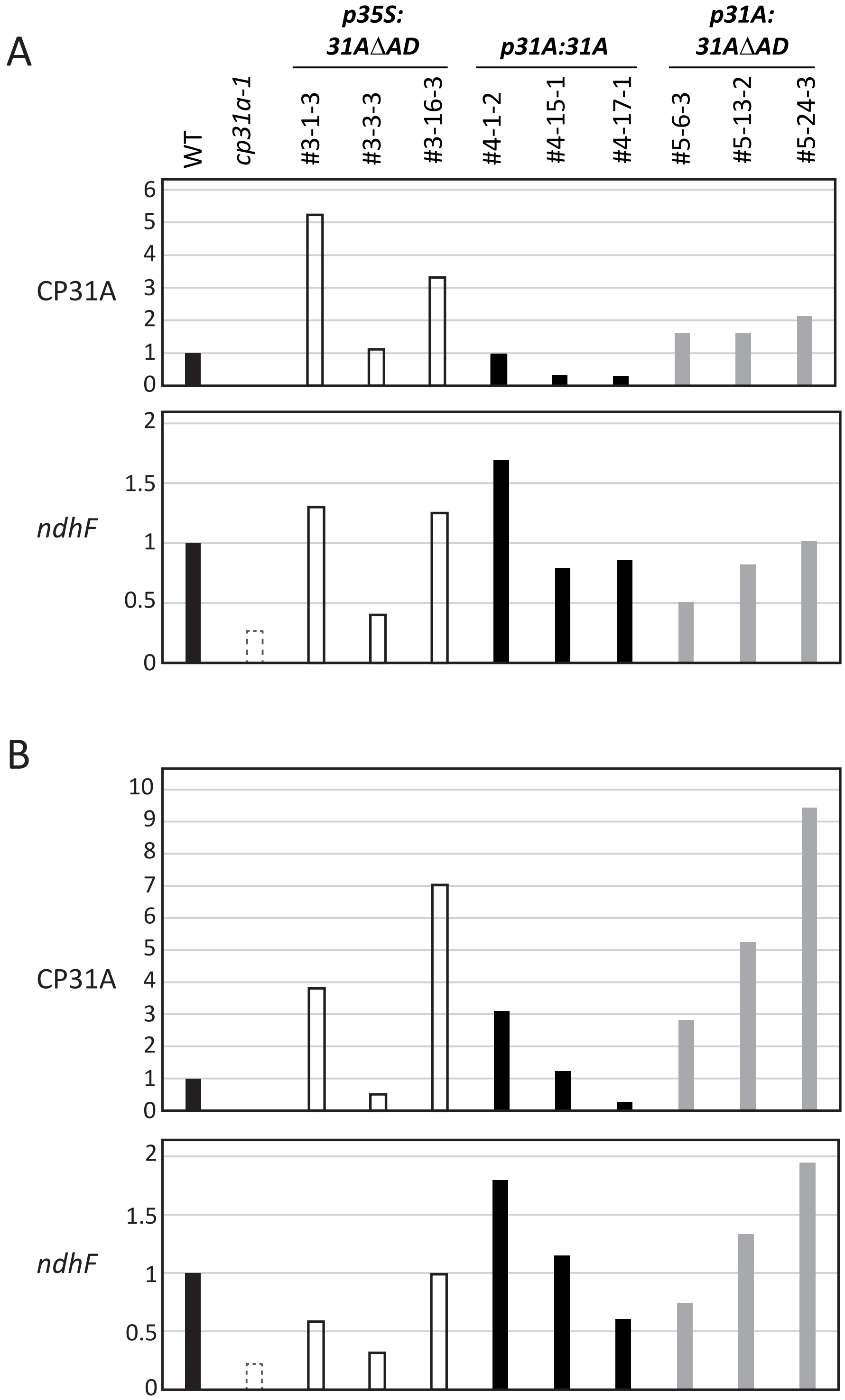


Supplemental Figure 2: Quantification of Immunoblots and RNA gel blot hybridizations shown in Fig. 4B and C.

A) Quantification of CP31A protein immunoblot signals and *ndhF* RNA gel blot hybridization signals shown in Fig. 4B. These values were used to calculate the *ndhF*/CP31A ratio shown in Fig. 4B.

B) Same as A, but with tissue only from the centre of the rosette (bleached area in *cp31a* null mutants, which corresponds to the data shown in Fig. 4C).

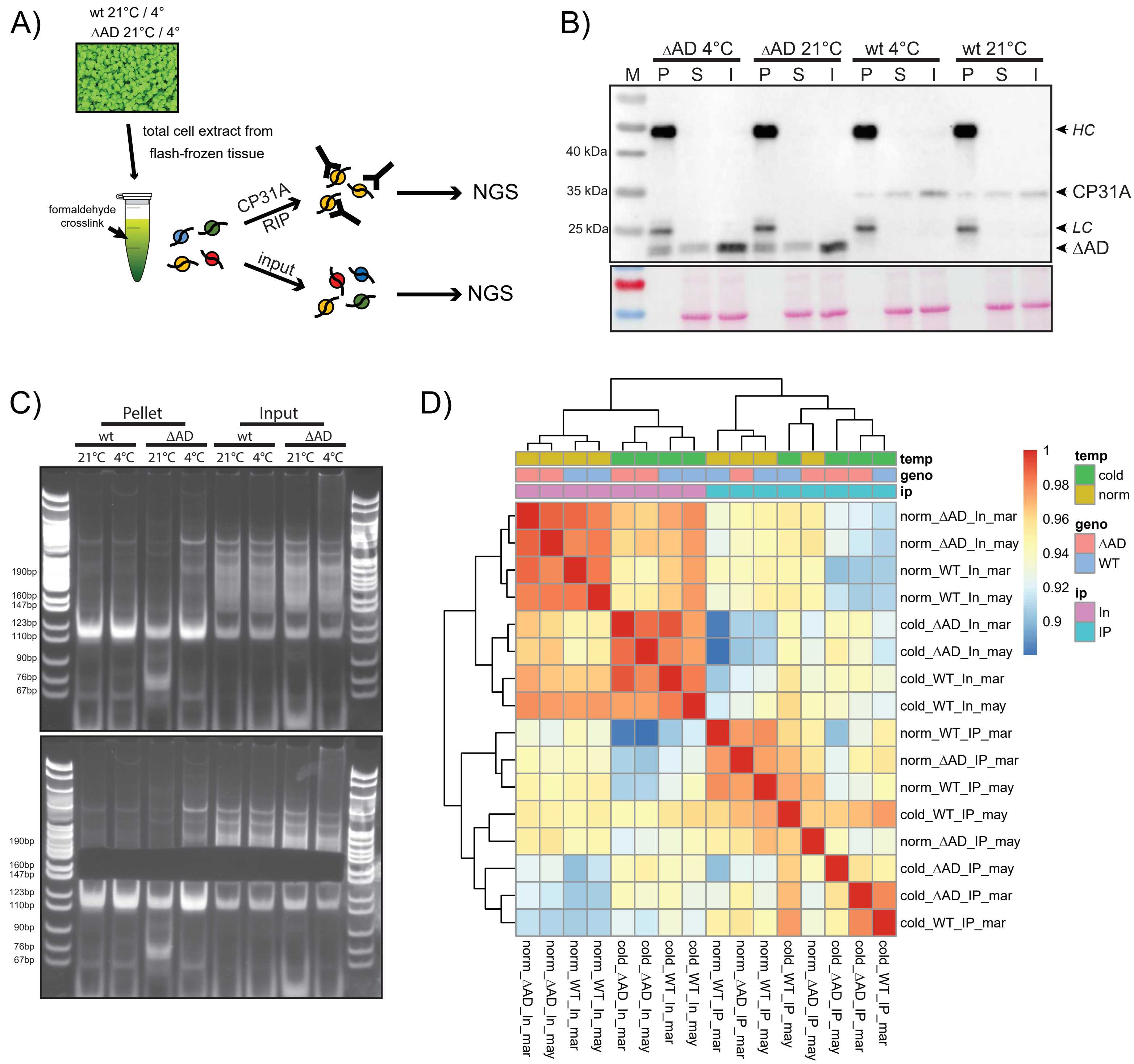


Supplemental Figure 3: RIP-seq analysis

A) Schematic overview of RIP-Seq. Seedlings are grown for 14 days prior to flash-freezing and whole cell lysate preparation. Lysates are cross-linked with formaldehyde and RNA binding proteins (here cpRNPs) are immunoprecipitated. RNAs from precipitates and from the input were analysed by next generation sequencing (NGS).

B) Immunoblots of RIP-fractions from wt plants or plants expressing a CP31A protein devoid of the acidic domain (-AD) grown either at 21°C or at 4°C. The blots were probed with the affinity-purified CP31A antibody used in the RIP-Seq assays. The upper panel shows the signals after chemiluminescence detection; the lower panel shows an excerpt of the corresponding Ponceau stain. IgGs in the immunoprecipitates (HC = heavy chain; LC = light chain) are detected by the secondary antibody used to probe the immunoblot. -AD = IPs from plants expressing CP31A without an acidic domain. P, S, and I = pellet, supernatant and input fractions of RIP experiments.

C) Purified RIP-seq libraries from pellet and input fractions were separated on a 6% Polyacrylamide gel for size-selection. The gel was stained with SYBR Gold nucleic acid gel stain and visualized on a UV transiluminator. Top: Gel prior to size selection. Bottom: Gel after size selection of all nucleotide bands between 160bp to 195bp. These bands correspond to adapter-ligated constructs derived from around 30 to 65 nucleotide RNA fragments. WT = *Arabidopsis thaliana* Col-0; -AD = AD deletion mutant.

D) Heatmap representing the Pearson correlation between the RIP-seq samples. Correlation was calculated over the DESeq2-normalized read counts over the candidate binding site regions.

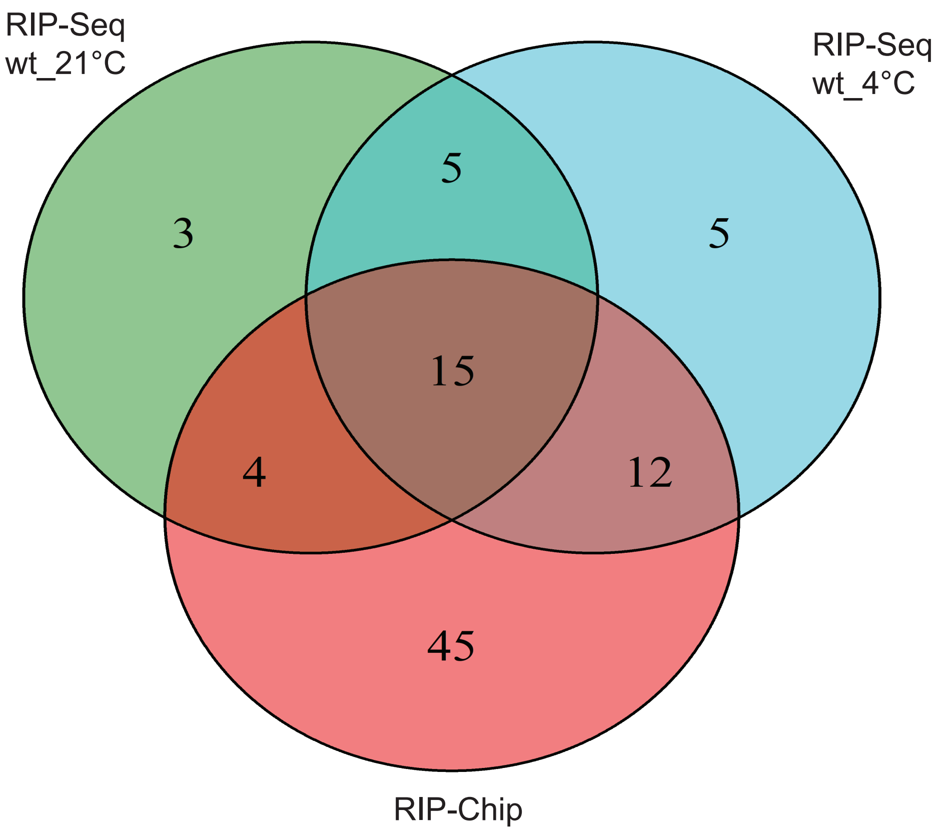


Supplemental Figure 4: Venn diagram summarizing the overlap of CP31A binding sites identified by RIP-Seq (WT_21°C and WT_4°C) with transcripts identified as CP31A-bound by previous RIP-Chip analysis (Kupsch et al. 2012). For the comparison significant CP31A binding sites (FDR < 0.05) were assigned to whole transcripts regardless of sense or antisense orientation. Binding sites that could not be assigned to a specific transcript and binding sites in transcripts not detected in the RIP-Chip (ndhB_2, rpl2_2, ycf2_2) were omitted. RIP-Chip probes with a signal above the median (median of ratios [635nm/532nm]) of all probes were considered as CP31A-bound. Probes spanning more than one gene were counted for all genes covered. RIP-Chip probes in intergenic regions were assigned to both adjacent genes.

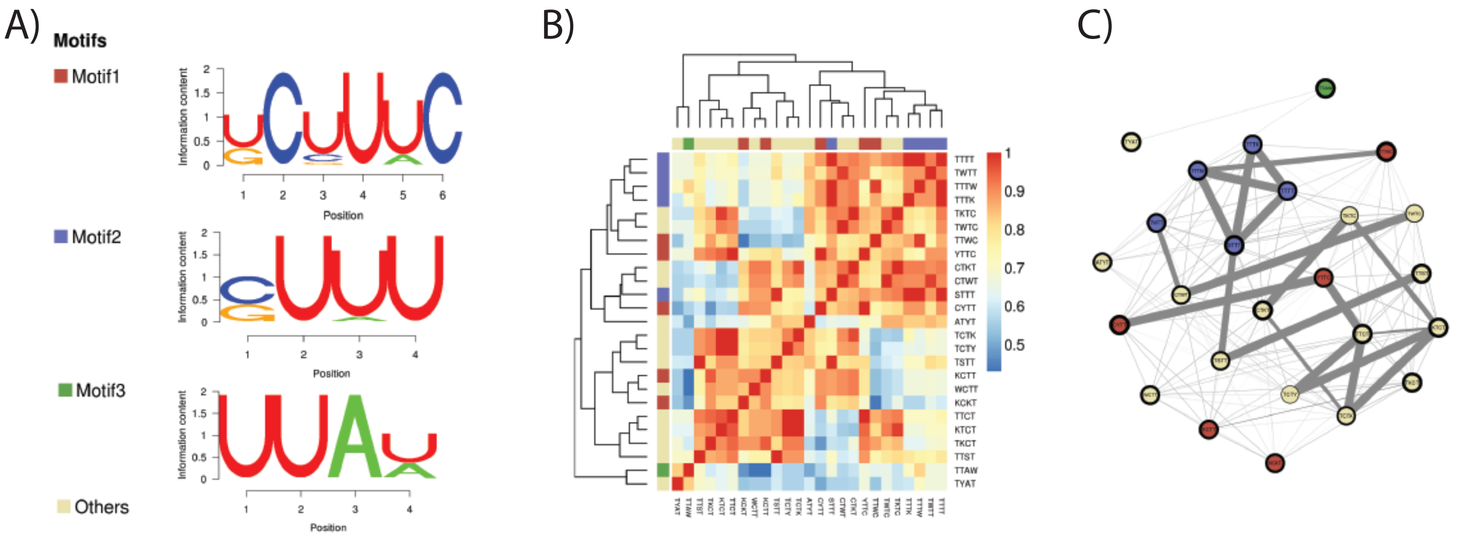


Supplemental Figure 5: Results of the SSMART RNA motif finder using CP31A BSs identified in wild type plants grown under cold conditions.
A) The three top scoring consensus sequence motifs from the SSMART search are shown. Each motif is assigned to colour which is used to indicte these motifs in (B) and (C).
B) Heatmap of the similarity scores between all unique evolved k-mers. The rows and columns represent k-mers and are color-coded to match the top 3 motifs. Together with (C), this forms the basis for the generation of consensus motifs by the SSMART algorithm.
C) Network graph of all unique evovled k-mers. The nodes represent the k-mers (the contour thickness correlates with its SSMART score). The edges represent the similarit score between k-mers: thin light fray lines correspond to a score > 0.7, while thick dark grey lines to a score > 0.9.

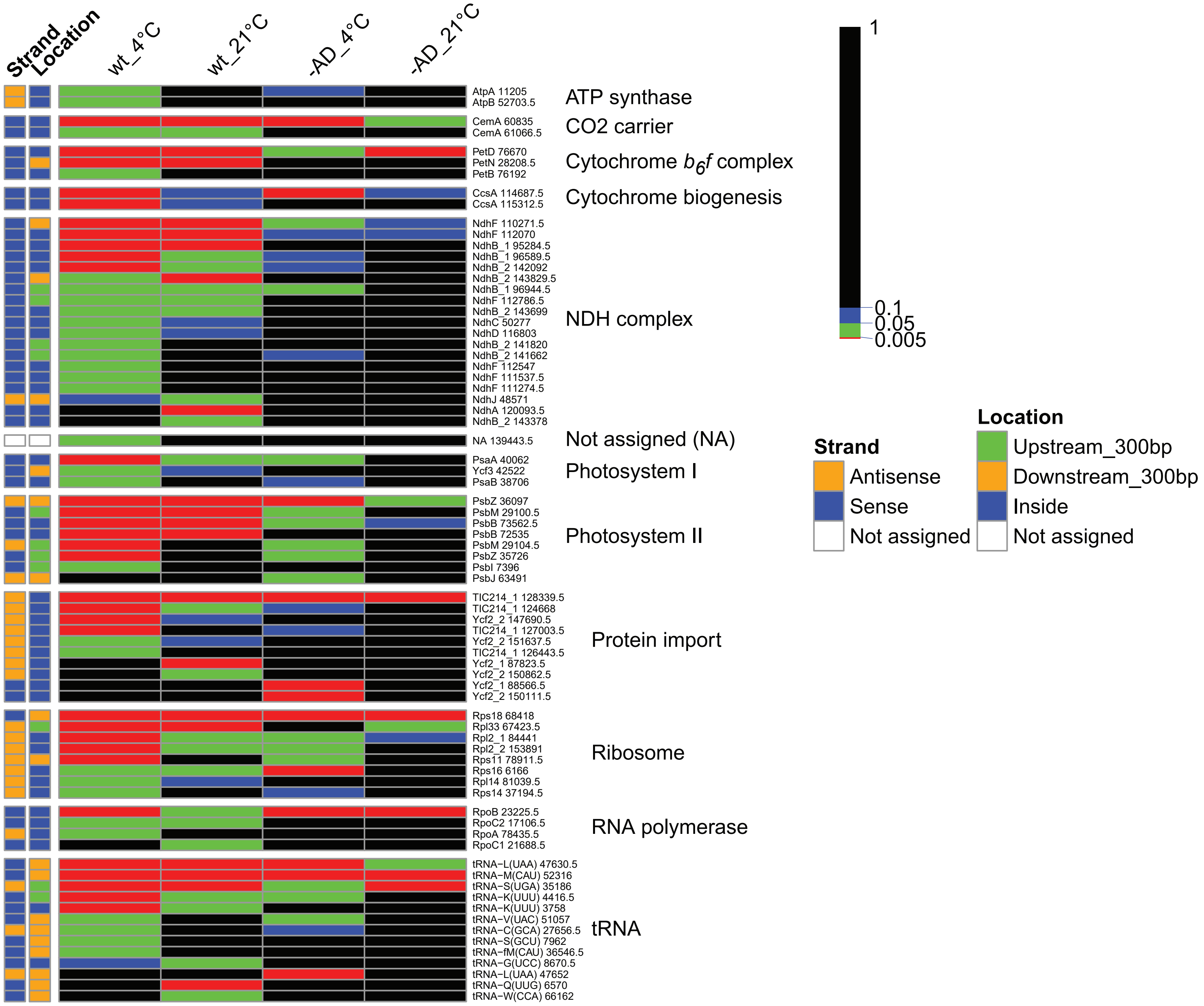


Supplemental Figure 6: Significance analysis of CP31A binding sites
Heatmap summarizing the levels of significance (FDR) for CP31A binding sites in the two genotypes and temperature conditions analyzed. Each binding site was assigned to a potential target gene, when the BSs location overlap the gene region and/or the adjacent +/- 300 bp (NA = no gene was near) and labelled with a number that refers to the middle position of the detected binding region. The genes are grouped according to their functional type. The legend left of the heat map indicates: (i) if the BS is targeting the sense or antisense transcript of the respective gene, and (ii) the location of the BS relative to the targeted gene.
