## Supplemental Tables for "Chloroplast cold-resistance is mediated by the acidic domain of the RNA binding protein CP31A"

Supplemental Table 1: Comparison of CP31A binding sites identified by RIP-Seq and transcripts associated with CP31A according to a previous RIP-Chip analysis (Kupsch et al. 2012).

| Gene[1] | Type | Binding Site WT cold (RIP-seq)[2] | Binding Site WT normal (RIP-seq)[2] | RIP-Chip (Kupsch et al. 2012)[3] |
|---|---|---|---|---|
| AtpA | ATP synthase | + | – | + |
| AtpB | ATP synthase | + | – | + |
| CcsA | cytochrome biogenesis | + | – | + |
| CemA | CO2 carrier | + | + | + |
| NdhA | NDH complex | – | + | + |
| NdhB | NDH complex | + | + | + |
| NdhC | NDH complex | + | – | + |
| NdhD | NDH complex | + | – | + |
| NdhF | NDH complex | + | + | + |
| NdhJ | NDH complex | – | + | + |
| PetB | Cytochrome b6f complex | + | – | + |
| PetD | Cytochrome b6f complex | + | + | + |
| PsaA | Photosystem I | + | + | + |
| PsaB | Photosystem I | + | – | + |
| PsbB | Photosystem II | + | + | + |
| PsbM | Photosystem II | + | + | + |
| PsbZ | Photosystem II | + | + | + |
| Rpl14 | Ribosome | + | – | + |
| Rpl2 | Ribosome | + | + | + |
| Rpl33 | Ribosome | + | + | + |
| RpoA | RNA polymerase | + | – | + |
| RpoB | RNA polymerase | + | + | + |
| RpoC1 | RNA polymerase | – | + | + |
| RpoC2 | RNA polymerase | + | + | + |
| Rps11 | Ribosome | + | – | + |
| Rps14 | Ribosome | + | – | + |
| Rps18 | Ribosome | + | + | + |
| TIC214 | protein import | + | + | + |
| tRNA-S(UGA) | tRNA | + | + | + |
| Ycf2 | protein import | – | + | + |
| Ycf3 | Photosystem I | + | – | + |
| NdhB_2 | NDH complex | + | + | – |
| PetN | Cytochrome b6f complex | + | + | – |
| PsbI | Photosystem II | + | – | – |
| Rpl2_2 | Ribosome | + | + | – |
| Rps16 | Ribosome | + | + | – |
| tRNA-C(GCA) | tRNA | + | – | – |
| tRNA-fM(CAU) | tRNA | + | – | – |
| tRNA-G(UCC) | tRNA | – | + | – |
| tRNA-K(UUU) | tRNA | + | + | – |
| tRNA-L(UAA) | tRNA | + | + | – |
| tRNA-M(CAU) | tRNA | + | + | – |
| tRNA-Q(UUG) | tRNA | – | + | – |
| tRNA-S(GCU) | tRNA | + | – | – |

| | | | | |
|---|---|---|---|---|
| tRNA-V(UAC) | tRNA | + | – | – |
| tRNA-W(CCA) | tRNA | – | + | – |
| Ycf2_2 | protein import | + | + | – |

[1] Binding sites were assigned to whole transcripts regardless of sense or antisense orientation. NA binding sites were not included. RIP-chip probes spanning more than one transcript were counted for all transcripts. Rip-chip probes in intergenic regions were also assigned to both adjacent genes.

[2] Significant binding site with an FDR $< 0.05$: +; binding site with an FDR $\geq 0.05$: -

[3] All probes with a median(median of ratios (635/532) CP31A) above the median (median of ratios (635/532) CP31A) of all probes were considered CP31A-bound: +; not CP31A-bound: -.
Genes highlighted grey are not comparable between the different methods as there is no information for these genes in the RIP-chip analysis.

Supplemental Table 2: Oligonucleotides

| Oligonucleotide | Sequence (5'>3') | Target(s) | Purpose | Reference |
|---|---|---|---|---|
| CP31A5'up-1694 for | GTTGGTGTGATTAGTATGTGGC | CP31A 5' UTR | vector construction | This study |
| CP31A-CDS5'_rev | GAGGTAACTATAGAAGAAGCCAT | CP31A 5' UTR | vector construction | This study |
| CP31A-5'-671_rev | CTTATAATGAGCAATGATAAC | CP31A 5' UTR | vector construction | This study |
| CP31A-5'-1013_for | GCTGGATTTAGAATCATTAG | CP31A 5' UTR | vector construction | This study |
| A31-minusAD_rev | ATCGGGATCCCTGGGCAACGAAGGAGACAAAGGG | CP31A | vector construction | This study |
| A31-minusAD_for | ATCGGGATCCGCCAAGCTTTTCGTCGGAAATTTG | CP31A | vector construction | This study |
| SmaI-31A F01 | ACCCGGGATGGCTTCTTCTATAGTTACC | CP31A | vector construction | This study |
| 31A-SmaI R01 | ACCCGGGTTAATATCCACGCCTTGGAG | CP31A | vector construction | This study |
| pGL-xhoxba_rev | CGATCTAGACTCGAGAAGCTTCACGCTGC | pGL vector | vector construction | This study |
| bar_rev_1 | GACTTCAGCAGGTGGGTGTAG | pGL vector | vector construction | This study |
| 31A_A-AD_test_for | ATGGCTTCTTCTATAGTTACCTCTAGC | CP31A | vector construction | This study |
| XhoI-31Apro_for | CTCTCGAGTTGGTGTGATTAGTATGTG | CP31A 5'UTR | vector construction | This study |
| pGL-XhoI_for | CACTCGAGTCTAGAGTCGACCTGCAGCCCG | pGL vector | vector construction | This study |
| pGL-XhoI_rev | GTCTCGAGAAGCTTCACGCTGCCGCAAGCAC | pGL vector | vector construction | This study |
| matK.AT.for | CGTTACCGGGTAAAAGATGC | matK | editing analysis | Tillich et al. 2009 |
| matK.AT.rev | AGCGGCGTATCCTTTGTTGC | matK | editing analysis | Tillich et al. 2009 |
| rpoB.AT.3for | GAGGTGGGTTCAGAAAAAGG | rpoB | editing analysis | Tillich et al. 2009 |
| rpoB.AT.3rev | TATCTGTCCTACATTCATGCG | rpoB | editing analysis | Tillich et al. 2009 |
| rpoB.AT.for | GAAAACCAGTAGGAATATGC | rpoB | editing analysis | Tillich et al. 2009 |
| rpoB1seq2 | TCCCCACCTACACAAGAAAATTG | rpoB | editing analysis | Tillich et al. 2009 |
| psbZ.AT.for | GCTTTCCAATTGGCAGTTTTTG | psbZ | editing analysis | Tillich et al. 2009 |
| psbZ.AT.rev | CCACCAAGAAGACTAATCCAATCC | psbZ | editing analysis | Tillich et al. 2009 |
| rps14.AT.for | TTATAGGGAGAAGAAGAGGC | rps14 | editing analysis | Tillich et al. 2009 |
| rps14.AT.rev | TACCAGCTTGATCTTGTTGC | rps14 | editing analysis | Tillich et al. 2009 |
| petL.AT.rev | ATTTCAATTGAAACTTAGGG | petL | editing analysis | Tillich et al. 2009 |
| petLseq | GGTAATTAACACGGTAAGGAACTATCG | petL | editing analysis | Tillich et al. 2009 |
| ndhBex1.rp | CCGATGGAGAGAAGAACCTATG | ndhB | editing analysis | Tillich et al. 2009 |
| ndhB1seq | TGAACCATATAGCCAAGAGAAACC | ndhB | editing analysis | Tillich et al. 2009 |
| CK_ndhB_ex1.for | CTGAGCAATCGCAATAATCG | ndhB | editing analysis | Kupsch et al. 2012 |
| At ndhB ex2 rev | CAAATGGTGGATATGCGAGC | ndhB | editing analysis | Kupsch et al. 2012 |

| | | | | |
|---|---|---|---|---|
| ndhF.AT.for | AAAACCTTCGCCGCATGTGG | ndhF | editing analysis | Tillich et al. 2009 |
| ndhF.AT.for | AAAACCTTCGCCGCATGTGG | ndhF | editing analysis | Tillich et al. 2009 |
| ndhF.AT.rev | GCATTCGCTGCAATAGGTCG | ndhF | editing analysis | Tillich et al. 2009 |
| ndhD.AT.for | CAAGCCTAATTCTATCATAACTCG | ndhD | editing analysis | Tillich et al. 2009 |
| ndhD.AT.rev | AAGTTTATATGGTTCGAACG | ndhD | editing analysis | Tillich et al. 2009 |

Supplemental Table 3: Sequenced and aligned Illumina reads for each RIP-seq library

| **Samples** | **Reads after trimming** | **Aligned to the whole *Arabidopsis* genome[1]** | **Aligned to the *Arabidopsis* chloroplast genome** | **Aligned to the whole *Arabidopsis* genome (after de-duplication)** | **Aligned to the *Arabidopsis* chloroplast genome (after de-duplication)** | **Aligned to the *Arabidopsis* chloroplast genome (after de-duplication in %)** |
|---|---|---|---|---|---|---|
| 21°C_WT_In_1 | 13738854 | 19703143 | 4086926 | 6602579 | 1230100 | 18.6 |
| 21°C_WT_In_2 | 3551118 | 4584772 | 1144044 | 2580316 | 524959 | 20.3 |
| 4°C_WT_In_1 | 12379796 | 16295656 | 5014941 | 5293791 | 1262483 | 23.8 |
| 4°C_WT_In_2 | 5670240 | 7942502 | 2048463 | 3976782 | 859073 | 21.6 |
| 21°C_DAD_In_1 | 13372609 | 17802580 | 3851478 | 6557615 | 1236664 | 18.9 |
| 21°C_DAD_In_2 | 11724622 | 14888994 | 3515171 | 4789701 | 881963 | 18.4 |
| 4°C_DAD_In_1 | 15096992 | 19688359 | 6278275 | 6546137 | 1645441 | 25.1 |
| 4°C_DAD_In_2 | 5262942 | 6887294 | 2067910 | 3098307 | 685503 | 22.1 |
| 21°C_WT_IP_1 | 4431219 | 5538627 | 1533281 | 1603180 | 375123 | 23.4 |
| 21°C_WT_IP_2 | 7140021 | 9601927 | 3006988 | 1765434 | 453555 | 25.7 |
| 4°C_WT_IP_1 | 7488555 | 8737397 | 3117471 | 1419271 | 425652 | 30.0 |
| 4°C_WT_IP_2 | 17486698 | 24059707 | 7462373 | 2671977 | 678102 | 25.4 |
| 21°C_DAD_IP_1 | 2684339 | 3220484 | 793058 | 1442051 | 311538 | 21.6 |
| 21°C_DAD_IP_2 | 5481601 | 6835783 | 1619002 | 901994 | 168802 | 18.7 |
| 4°C_DAD_IP_1 | 6595978 | 8107400 | 2816224 | 2435052 | 754216 | 31.0 |
| 4°C_DAD_IP_2 | 1457784 | 1830217 | 641175 | 652789 | 176142 | 27.0 |

[1] more reads were aligned than sequenced because some reads align at several locations (e.g. inverted repeat region)
